# Proposed allosteric inhibitors bind to the ATP site of CK2α

**DOI:** 10.1101/2020.07.07.191353

**Authors:** Paul Brear, Darby Ball, Katherine Stott, Sheena D’Arcy, Marko Hyvönen

## Abstract

CK2α is a ubiquitous, well-studied protein kinase that is a target for small molecule inhibition, for treatment of cancers. While many different classes of ATP-competitive inhibitors have been described for CK2α, they tend to suffer from significant off-target activity and new approaches are needed. A series of inhibitors of CK2α has recently been described as allosteric, acting at a previously unidentified binding site. Given the similarity of these inhibitors to known ATP-competitive inhibitors, we have investigated these further. In our thorough structural and biophysical analyses, we have found no evidence that these inhibitors bind to the proposed allosteric site. Rather, we report crystal structures, competitive ITC and NMR, HDX mass spectrometry and chemoinformatic analyses that all point to these compounds binding in the ATP pocket. Our crystal structures however do show that the proposed allosteric site can bind ligands, just not those in the previously described series. Comparison of our results and experimental details with the data presented in the original report suggest several reasons for the disparity in our conclusions, the primary reason being non-specific inhibition by aggregation.

**Table of Content graphics:** 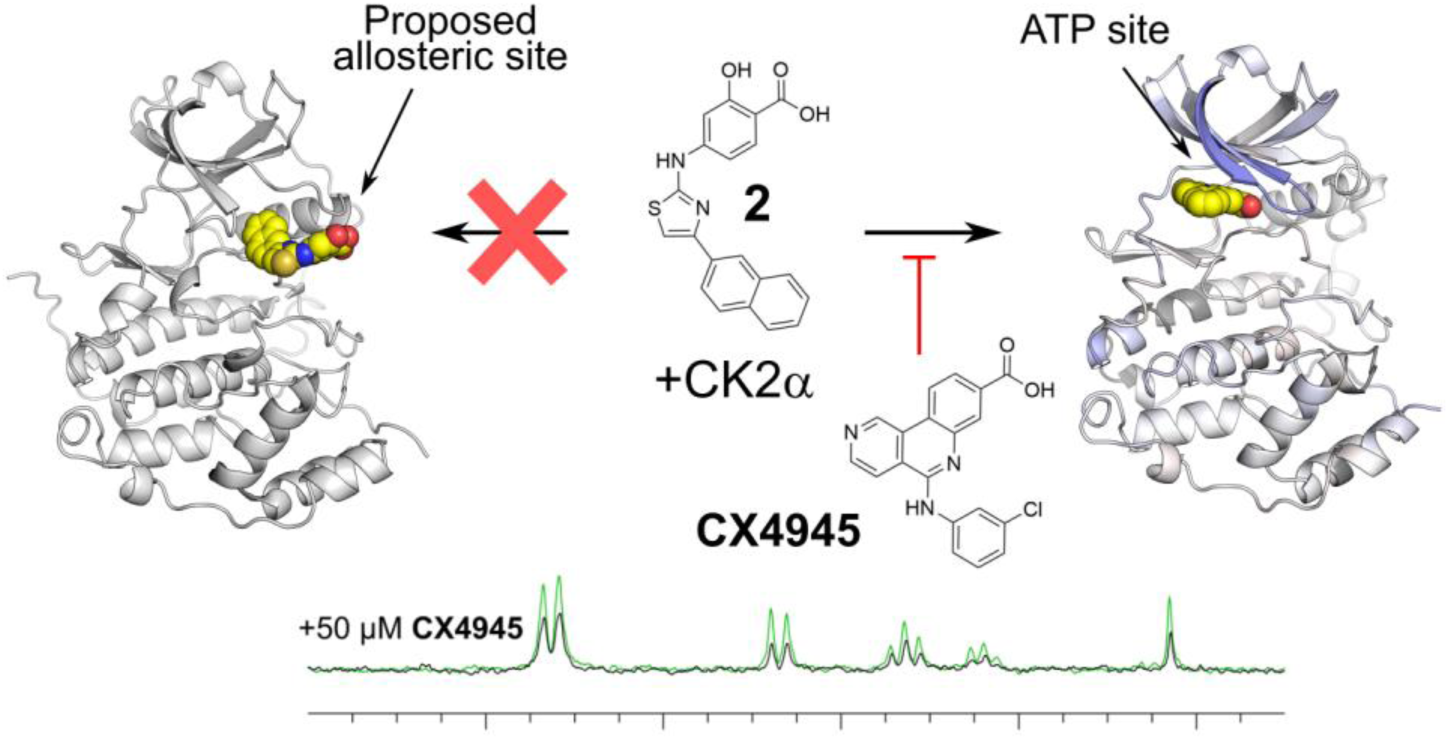

## Introduction

Potent inhibitors of kinases have existed for some time and the majority of these bind in the active site of the kinase, competing with the co-factor ATP^1,2^. Because of the significant conservation of the ATP binding site, kinase inhibitors often suffer from poor selectivity^3–7^. This poor selectivity is a disadvantage for several reasons. Firstly, promiscuous inhibitors tend to manifest off-target toxicity if used therapeutically and, to combat this, end up being used at suboptimal concentrations^8,9^. Secondly, they are poor chemical tools as they cannot provide unambiguous answers to target validation questions. In order to develop more selective inhibitors of kinases, a promising strategy is to target sites outside the highly conserved ATP pockets. This leads to increased selectivity as these sites are structurally not constrained by the shared property of binding to ATP and therefore tend to be less conserved among different kinases^10–13^. When developing inhibitors that bind outside the primary ATP site, it is vital that the new sites are thoroughly validated biochemically, biophysically, and structurally to confirm the inhibitor binding mode unambiguously. Often inhibitors are identified as being allosteric if they are shown to be non-competitive in biochemical assays. However, this data can be misinterpreted, as non-specific inhibitors also often show similar non-competitive behaviour^14^. This problem can be further compounded by moderate/weak affinity of the hits being investigated, as they must be used at higher concentrations, which makes secondary binding more likely. Therefore, apparent allosteric behaviour must be very carefully validated to ensure that vital time and resources are not wasted on optimising these inhibitors further.

CK2α has a long history as a target for drug discovery, due to its ubiquitous role in multiple diseases such as cancer and fibrosis^15–17^. CK2α is an unusual kinase in that it is constitutively active and does not require phosphorylation for activation^18^. It is also a highly promiscuous kinase with hundreds of recorded cellular substrates.^19^ Inhibition of CK2α has been achieved mostly through an ATP-competitive mechanism and a number of high affinity inhibitors of distinctly different chemotypes are known and characterized structurally^17,20–27^. Despite significant effort, most inhibitors have poor selectivity, although claims are often made to the contrary^28–32^. For example, the clinical trial candidate CX-4945 inhibits at least ten other kinases with nanomolar affinity (see Table S1 in Brear et al.^25^).

We have recently demonstrated a new approach to CK2α inhibition. By using a cryptic so-called αD pocket just below the active site as an anchoring point, we have been able to develop highly selective inhibitors of CK2α^25^. The most potent of these is the most specific CK2α inhibitor reported to date, CAM4066^25^. The fragment-based development of these inhibitors was supported by a significant amount of structural data to identify the optimal αD site anchor moiety and subsequent growth of the fragment to the ATP site^27^. In addition to active site targeting, another approach is the inhibition of CK2α binding to its scaffolding partner CK2β. This method has the potential to generate specific inhibitors that affect only a subset of CK2 substrates^24,33^ and has led to some initial encouraging success that requires further development^24,33,34^.

Recently, a series of CK2α inhibitors (Series A, SI Fig. 1), proposed to bind in a novel site (Site A, SI Fig. 2) just outside of the highly conserved ATP site, has been published^35,36^. The authors used enzymatic assays, native mass spectrometry and competitive ligand-based NMR studies to characterize the mode of inhibition. They interpret their data as supporting an allosteric mechanism. While their data were presented as supporting non-ATP-competitive inhibition, there were several reasons why we speculated that the mechanism of inhibition by these molecules was not necessarily allosteric. In particular: the enzyme assays showed mixed mode inhibition; ATP analogues were replaced in NMR studies; high affinity ligands showed unexpected signal in ligand-based NMR experiments; and, mutations close to the ATP site affected inhibitory potential ^35,36^. Our motivation to investigate these inhibitors in more detail arose from the somewhat conflicting data presented, ^35,36^ as well as the chemical properties of these new inhibitors, which are not unlike known ATP competitive kinase inhibitors.

**Figure 1.**
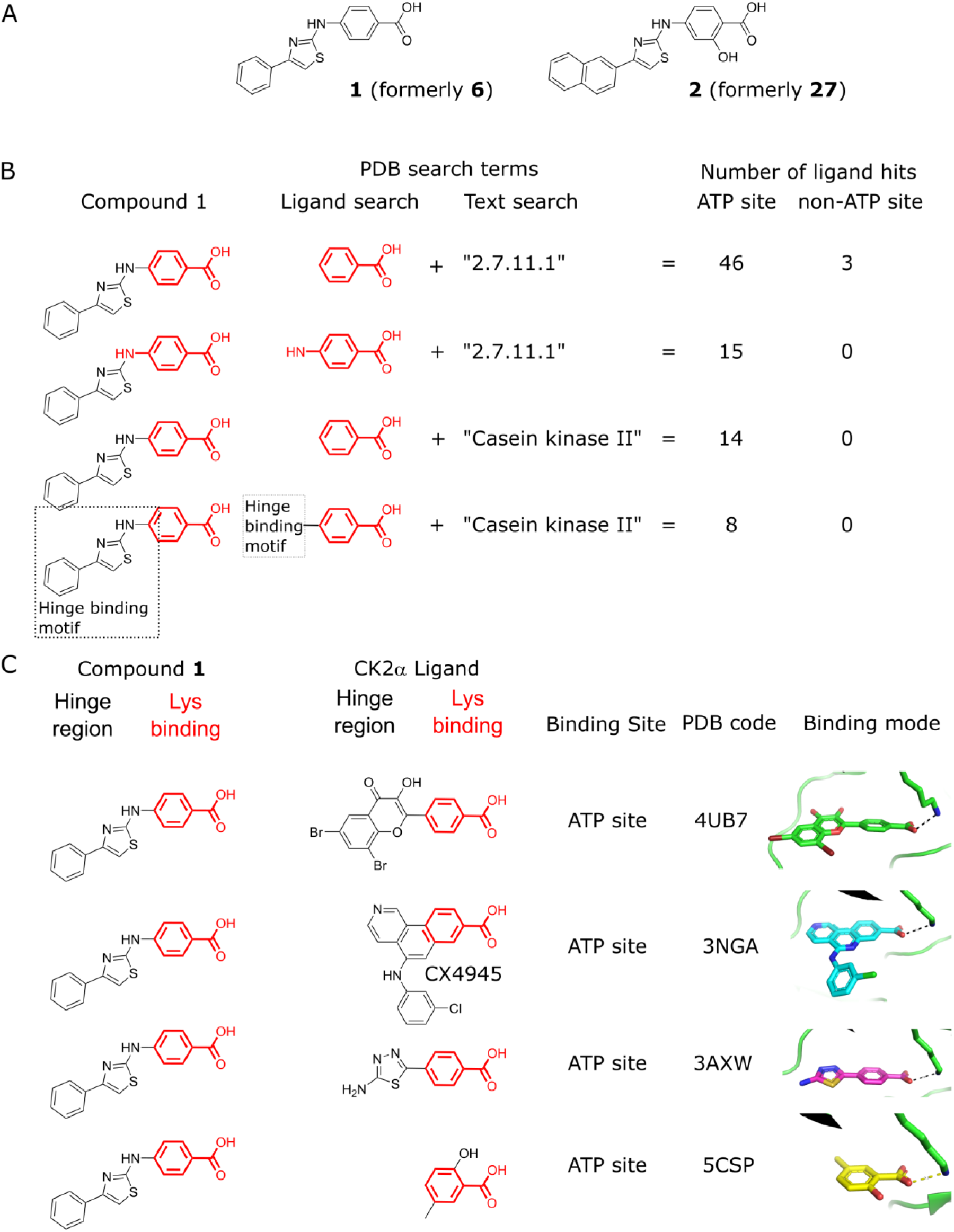
Conserved pharmacophore binding in the ATP site of protein kinases. (A) The structures of **1** and **2**, denoted **6** and **27** respectively in the original studies. (B) A summary of the analysis of the established ligands that bind to Ser/Thr protein kinases (EC number 2.7.11.1) or CK2α in the ATP site. (C) The structures of the four CK2α inhibitors and their protein-bound crystal structures which contain a benzoic acid group that binds in the ATP site^27,37,46–55,38,56–63,39–45^.

**Figure 2.**
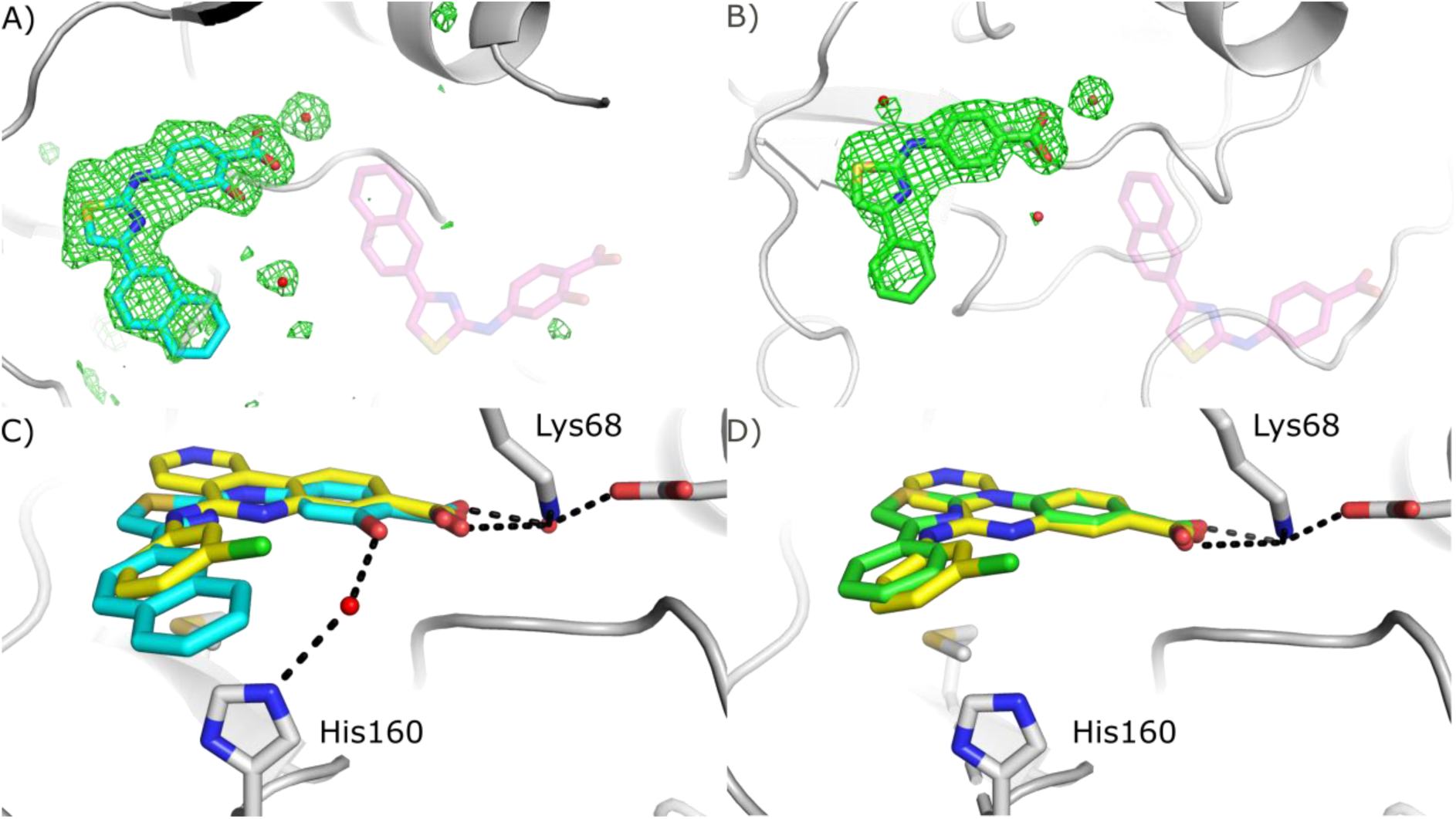
Crystal structures of compounds 1 and 2 bound to CK2α. (A) The Fo-Fc map (green, displayed at 3s) in the ATP site and Site A before the ligand has been placed. Refined structure of **1** (green) has been superimposed on the ATP site (PDB: 6YPJ) and the predicted binding mode of **2** (transparent purple) is shown on Site A. (B) The Fo-Fc map (green, contoured at 3 s) for the ATP site and Site A before the ligand has been placed. Refined structure of **2** (blue) has been superimposed on the ATP site (PDB: 6YPG) and the predicted binding mode of **2** (transparent purple, from supplementary material of reference 34) is shown on Site A. (C) **1** (green) bound to CK2α, (PDB: 6YPJ) with key interacting residues and H-bonding highlighted. (D) **2** bound to CK2α (PDB: 6YPG), with key interacting residues and H-bonding highlighted. The structure of **CX4945** (PDB:3PE1, yellow) has been superimposed on the structure of **1** and **2** in panels C and D. Electron densities of the final, refined structures are shown in SI Fig. 6.

We have used several orthogonal biophysical and structural methods for the characterization of the binding mode of these new CK2α inhibitors and we find no evidence for interaction with the proposed allosteric site (Site A). We show, using crystal structures, competitive ITC and NMR, hydrogen-deuterium exchange mass spectrometry (HDX) and computational analyses of the ligand structures, that these molecules are clearly binding to the ATP site of the kinase. It thus seems likely that most of the inhibitory effect, if not all, comes from traditional type I inhibition of the kinase through orthosteric competition with ATP.

## Results

### Chemoinformatic analysis

The structures of the compounds presented in the original papers and their comparison to other validated kinase and CK2α inhibitors raised concerns over the claim that these compounds had a novel binding mode. The conserved scaffold of the compounds shares significant similarity to many established and well validated kinase inhibitors that bind in the ATP site (Fig. 1)^27,37–63^. This scaffold consists of a central thiazole ring that is connected directly to a phenyl group on one side and to a benzoic acid moiety through an amine on the other side (Fig. 1A). We have used two representative compounds from Series A for further investigation, which we denote **1** and **2** (these were referred to as **6** and **27** respectively in the original papers^35,36^). **1** was chosen as much of the validation of Series A and Site A relied upon this compound^35^. **2** was chosen as it was the highest affinity inhibitor reported in Series A.^36^ A search of the Protein Data Bank (PDB, http://www.pdb.org) for ligands that bind to serine/threonine kinases (EC=2.7.11.1) and contain a benzoic acid group, which is a critical part of the inhibitors in question (Fig. 1B), identified 49 molecules. Of these, 46 bind in the ATP site and the remaining three bind in interface sites that do not exist in CK2α (SI Table 1). Narrowing the search by inclusion of the amine that links the benzoic acid to the thiazole ring identified 15 ligands (SI Table 2). Of these ligands, all 15 bind in the ATP site. When a similar search was performed on CK2α, 14 ligands were identified, all of which bound in the ATP site (SI Table 3). Four representative structures of these compounds are shown in Fig. 1C. All of these inhibitors display the same binding mode where the binding to CK2α is dominated by the interaction between the carboxylic acid and the ζ-amino group of Lys68 (Fig. 1C). Further refining the search to include only compounds that also bind in the hinge region, similar in structure to the thiazole-amine group of **1**, identified eight inhibitors (SI Table 4). These compounds, including **CX4945**, an extensively validated ATP-competitive CK2α inhibitor currently in clinical trials, interact with the hinge and Lys68^47^. Overlaying the 2D structure of **1** with **CX4945** shows that **1** could replicate the binding mode of **CX4945** (SI Fig 2) and bind in the ATP site with its carboxylic acid interacting with Lys68 and amine hydrogen bonding with the hinge (Fig 1C). Indeed, modelling of **1** into the ATP site of CK2α replicates the crystallographic binding mode of **CX4945** (SI Fig 2). The carboxylic acid is interacting with Lys68 and an ATP site water as predicted and the hydrophobic core of the molecule is sandwiched between Met163 and Val66. The amine is not predicted to interact with the hinge. However, the benzyl group is predicted to bind in a similar position to the benzyl chloride of **CX4945**.

Further evidence in support of our hypothesis that **1** may bind in the ATP site comes from the analysis of a fragment-based X-ray crystallographic screen against CK2α^27^ previously performed at our lab. One purpose of fragment-based X-ray crystallographic screening is to use the fragments to probe the target protein and identify binding hotspots^64^. From the screen of 354 fragments, 21 were identified as binding to CK2α. All 21 bound in the ATP site and interacted with Lys68^27^. 19 of the CK2α-binding fragments were structurally related to **1** and contained a carboxylic acid moiety that mediated this interaction (SI Fig. 3). The preference of CK2α for binding ligands with a carboxylic acid in this position is further supported by the serendipitous binding of acetate. Acetate ions can be utilised as a very low molecular weight probe to identify binding sites for carboxylic acids^64^. In CK2α structures deposited to the PDB, at least nine binding sites for acetate can be identified (SI Fig. 4). Among these sites are the Lys68 hotspot in the ATP site, the CK2β interface and the substrate-binding pocket. Notably, none of the nine sites overlaps with Site A, the proposed binding site of **1**. When such a conserved binding mode is observed for an ATP site-binding pharmacophore, it seems unlikely that inhibitors with significant structural similarity would not also bind to the ATP site. These observations led us to initiate additional studies to investigate the binding mode of these inhibitors using two compounds (**1** and **2**, Fig. 1A) from the original publications.

**Figure 3.**
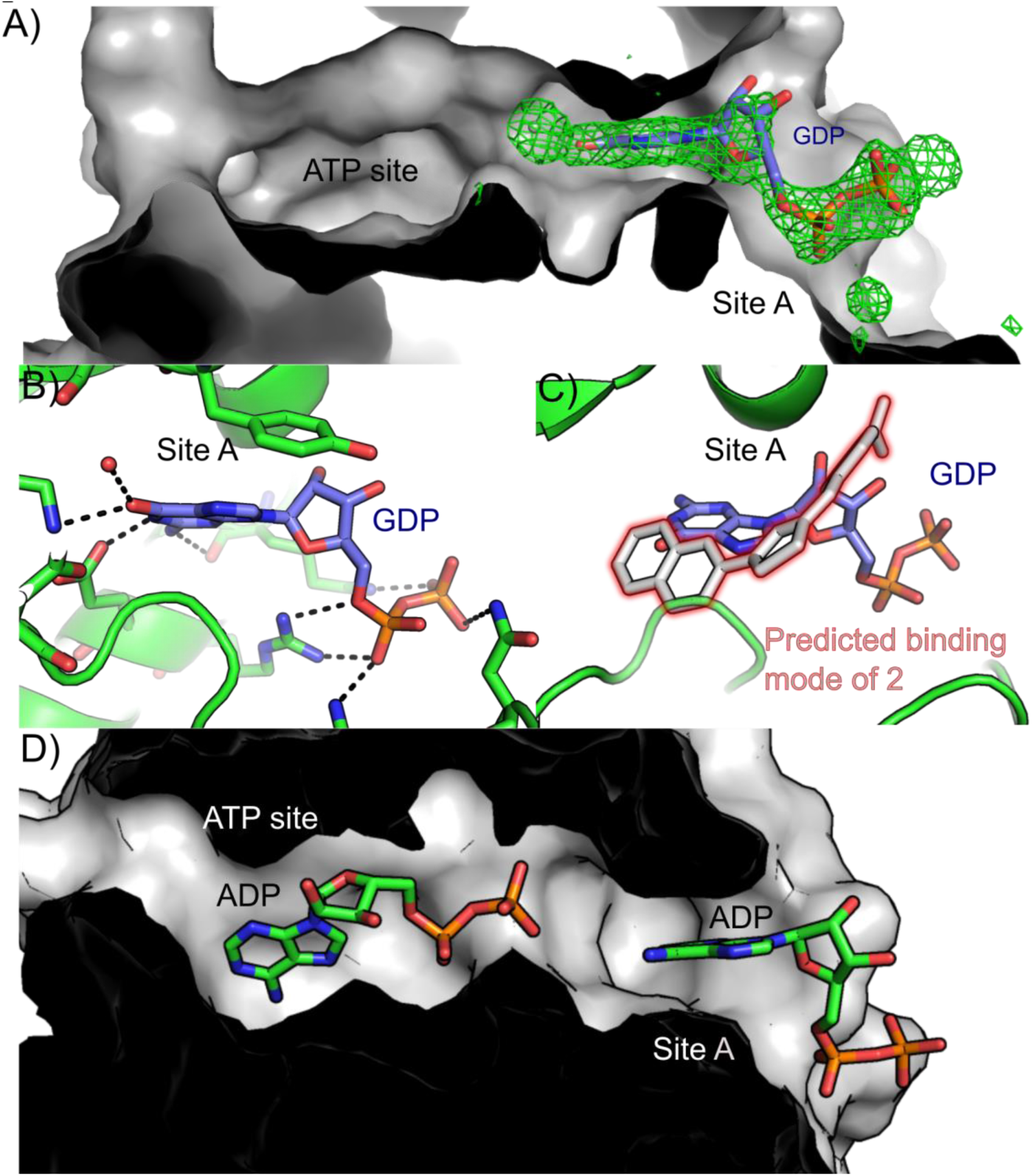
Crystal structures of ADP and GDP binding to CK2α Site A. (A) GDP binding in Site A. The Fo-Fc map (contoured at 3s) centered on the proposed allosteric site. No density was observed in the ATP site for GDP therefore the density in the ATP site was not shown for clarity. The CK2α KKK/AAA construct was used for this structure (PDB: 6YPK). B) The structure of GDP bound to CK2α, with the extensive H-bonding network shown by dotted lines (PDB: 6YPK). C) The proposed binding mode of **1** (PDB: 6YPJ) superimposed on the structure of GDP bound to CK2α (6PYK). D) The structure of ADP bound simultaneously in the ATP site and in Site A of WT CK2α (PDB: 6YPN).

**Figure 4.**
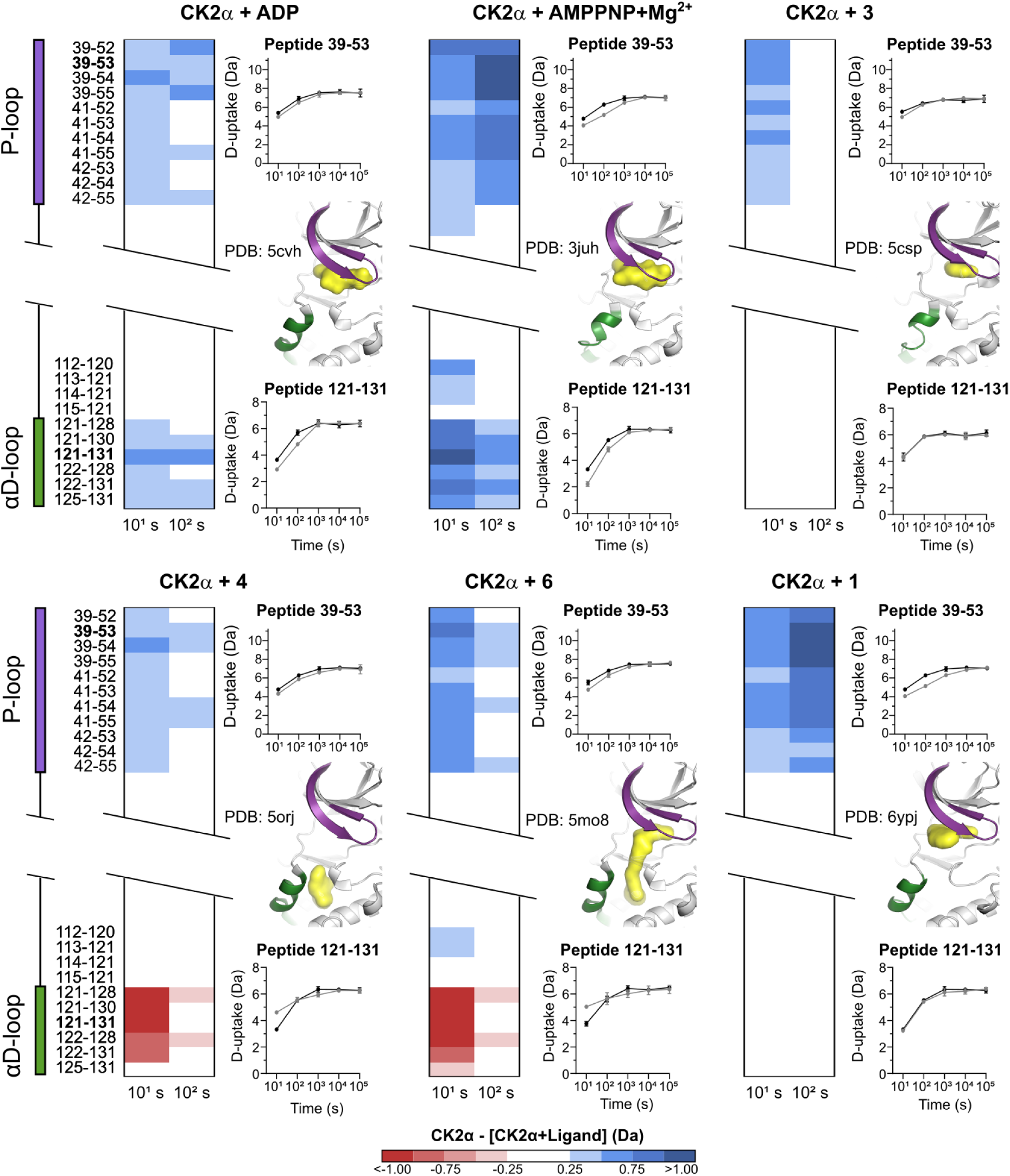
HDX analysis of ligand binding to CK2α. Heatmaps show the change in deuterium uptake upon binding of different ligands to apo CK2α. Each panel, with the ligand listed on the top, shows the heatmaps for 10 s and 100 s time points for peptides corresponding to the P-loop and αD-loop, in comparison to unliganded CK2α. All depicted changes are greater than ±0.25 Da and have a p-value less than 0.01 in a Welch’s t-test. The exact start and end residues for each peptide are shown on the left of the panels. Next to each heatmap are two graphs showing deuterium uptake for one peptide from each P-loop (top) and αD loop (bottom) for apo CK2α (black) and CK2α with ligand (grey). The Y-axis range is 80% of max D-uptake assuming the N-terminal residue undergoes 100% back-exchange. Data have not been corrected for back-exchange. Error bars are ±2s derived from three technical replicates. A structural diagram in between the uptake plots has the P-loop and αD loop coloured purple and green, respectively, and the bound ligand in surface rendering in yellow. Full HDX data are shown in SI Fig. 11 and available in the supplementary spreadsheet.

### X-ray crystallographic analysis

The original publication contained no structural data on the inhibitor binding mode. In fact, the authors state that ‘cocrystallisation attempts aimed at elucidating the non-ATP competitive binding mode of 2-aminothiazole derivatives with CK2α were not successful’^35,36^. We have rectified this and have determined structures of both **1** and **2** in complex with CK2α (Fig. 2, SI Table 5). These structures were obtained by soaking the compounds into CK2α crystals at 10 mM for 16 hours. The crystals diffracted at less than 2 Å resolution and unambiguous positive difference electron density corresponding to both **1** and **2** can be clearly seen in the ATP site before the ligands were included (Fig. 2A and 2B, SI Fig. 5). As we predicted, the carboxylic acid groups of both compounds interact directly with Lys68 like the carboxylic acid of **CX4945** (Fig. 2C and 2D). The remainder of the molecules however, each have their own binding nuances. The amine of the benzylamine of **1** interacts very weakly with the hinge region backbone nitrogen of Val116 via a bridging water molecule whereas the OH group of **2** interacts with His160, via a bridging water, which pulls the amine of the benzylamine further away from the hinge region and prevents interactions with the hinge (Fig. 2C and 2D). The 5-membered rings from **1** and **2** stack on top of Met163 which leads to the naphthyl ring of **2** stacking on top of His160 at the entrance to the ATP site. The observed binding mode agrees well with the binding mode of established ATP site inhibitors of CK2α, such as **CX4945**. No density corresponding to either **1** or **2** was observed in the proposed allosteric binding site (Site A, SI Fig. 5), despite the high concentration of the ligand (>1,000-fold excess over K_D_) used for soaking. The binding mode of **1** in the ATP site of CK2α observed in the crystal structure agrees well with that predicted by the modelling study (SI Fig. XXXX).

**Figure 5.**
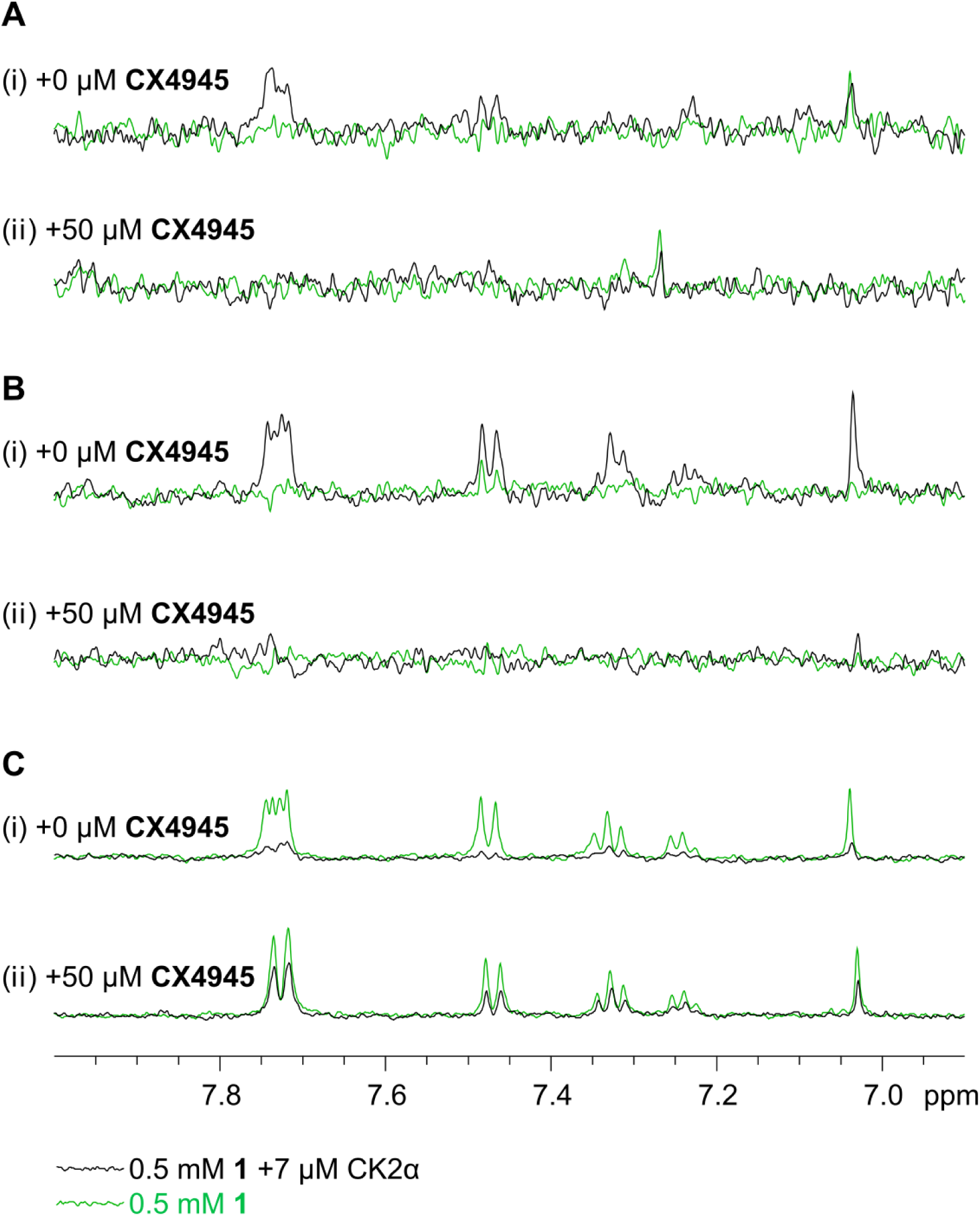
Ligand-based NMR experiments to detect binding of 1 to CK2α and competition by CX4945. All panels are overlays of spectra recorded in the presence (black) and absence (green) of CK2α. (A) Saturation Transfer Difference (STD) ^1^H spectra of **1** (i) in the absence of **CX4945**, and (ii) in the presence of 50 μM **CX4945**. (B) WaterLOGSY spectra of **1** (i) in the absence of **CX4945**, and (ii) in the presence of 50 μM **CX4945** (ii). (C) Relaxation-edited (CPMG) ^1^H spectra of **1** (i) in the absence of **CX4945**, and (ii) in the presence of 50 μM **CX4945** (ii). (NB: the signals at ca. 7.7 ppm are in fact two overlapping doublets and therefore small changes to either chemical shift results in an apparent change to the multiplet structure.)

To reduce the impact that different crystal forms and packing may have on the accessibility of binding sites and bias in the resulting complex structure, we have soaked the inhibitors into two different crystal forms of CK2α. One is obtained using an engineered form of CK2α (referred to as CK2α KKK/AAA) in which lysines 74-76 are mutated to alanines to facilitate crystallization. These lysines are in the vicinity of Site A and could affect binding of ligands to this site. However, these lysines do not affect the ATP site and we show later that both **1** and **2** bind to CK2α KKK/AAA and to wild type CK2α with the same affinity (see ITC measurements). The second crystal form (CK2α SF) has no mutations in the lysine-rich loop but has a substrate peptide fused to the N-terminus of the protein, aimed at facilitating crystallization. In both crystal forms, we observe electron density for **1** and **2** only in the ATP site, and not in the proposed allosteric site (Site A, SI Fig. 5).

The two crystal forms of CK2α have been extensively used for the determination of CK2α:ligand complexes (66 structures currently in the PDB). More than eight different ligand binding sites can be accessed by soaking into these crystal forms (SI Fig. 7), including the proposed allosteric site for **1** and **2**, Site A. We have observed both ADP and GDP bound to Site A in crystal structures of wild type CK2α (PDB:6YPN) and in the KKK/AAA mutant (PDB:6YPK) (Fig. 3). The adenosine and guanosine rings sit in the hydrophobic pocket identified in the previous modelling work^35^ and the diphosphate interacts with Arg80 that has been identified as an acetate-binding hotspot (SI Fig. 4). This confirms that Site A is accessible to ligands in these crystal forms and soaking with **1** or **2** would be expected to result in clear electron density in the site, should they bind there. Indeed, ADP can be seen simultaneously in both the ATP site and in Site A (Fig. 3D, PDB: 6YPN). These observations demonstrate that binding in Site A does not in itself prevent binding in the ATP site and that if **1** or **2** did bind in both sites, we would expect to see density corresponding to it. Furthermore, **1** and **2** would be expected to bind predominantly in the highest affinity site and this site would dominate the inhibition. These crystal structures prompted us to perform further experiments to unambiguously establish the binding site in solution and verify the likely mode of inhibition for these compounds.

### Hydrogen-deuterium exchange mass spectrometry

Hydrogen-deuterium exchange (HDX) allows the localization of interaction sites and conformational changes in proteins. We have analysed the binding of four different classes of ligands to CK2α (SI Fig. 8 and 9) using HDX. These ligands were: ADP and AMPPNP-Mg^2+^, both of which bind across the entire ATP site interacting with the hinge region and Lys68 (SI Fig. 8C and 8D);^60^ **3**, which is a fragment that binds only on the right hand side of the ATP site, interacting with Lys68^60^ (SI Fig. 8E); **4**, which is an αD site binder (SI Fig. 8G)^26^ and **6** which links the αD site and the ATP site (SI Fig. 8F)^27^. We recovered 192 shared peptides from each of these HDX experiments that redundantly span almost the entire sequence of CK2α (SI Fig. 10). This allows for unbiased detection of ligand interaction fingerprints across the whole of the protein. Binding can be identified by observing changes in deuterium uptake between CK2α alone and a concurrently run ligand-bound sample (SI Table 6). We show all changes greater than ±0.25 Da that have a p-value less than 0.01 in a Welch’s t-test (Fig. 4 and SI Fig. 11). Extended data for all peptides and timepoints are shown in the Supplementary Information (SI Fig. 11 and a separate spreadsheet). For all the ligands, a decrease in deuterium uptake is observed at late timepoints particularly for peptides containing residues 54-111 (sample peptides shown in SI Fig. 12). Since this change occurs with all the ligands, it appears to result from general stabilization of the region termed the ‘N-lobe’, on ligand binding, rather than an effect attributable to a specific site.

Addition of ADP and AMPPNP-Mg^2+^ causes clear changes in deuterium uptake in the ATP site (Fig. 4). The largest change is a decrease in deuterium uptake for peptides covering residues 39-55 and 121-131. Residues 39-55 correspond to the P-loop that interacts directly with the phosphates of ADP, while residues 121-131 are the αD loop that does not interact directly with ADP or AMPPNP-Mg^2+^. Protection occurs in the latter due to the interaction of the adenosine ring with the hinge region that directly links to and stabilizes the αD loop. These observations are similar to those previously observed when binding of ATP analogues to other kinases has been studied^65,66^. The observed decreases in uptake are larger for AMPPNP-Mg^2+^ than ADP-Mg^2+^ because AMPPNP binds with a higher affinity than ADP. To further probe interactions in a more localized area of the ATP site, we used **3** that binds only at the right hand side of the ATP site (SI fig 8 E), coordinated to catalytic Lys68 (PDB: 5CSP)^60^ (Fig. 4). With **3**, we see a decrease in deuterium uptake in the P-loop and no change in the αD loop. The pattern of protection within the P-loop is also distinct, indicative of different binding contacts being used by **3** compared to the ATP analogues. We can thus distinguish binding in the entire ATP site, including interaction with the hinge, from binding on just one side where Lys68 is located.

To ensure we could detect different binding sites with our HDX experiments, we used another well characterized ligand (**4**) that binds to the αD pocket only (SI Fig. 8G)^26^. This compound shows a markedly different HDX fingerprint compared to the ATP analogues and **3** (Fig. 4). The largest differences are seen in the αD loop, with binding of **4** resulting in increased uptake compared to unbound CK2α. Increased uptake in the αD loop is opposite to the decrease seen upon ADP and AMPPNP-Mg^2+^ binding. This indicates that the binding of **4** in the αD pocket puts the loop in a more flexible or open conformation amenable to exchange, as is seen in the crystal structure of CK2α in complex with **4** (PDB: 5ORJ). Finally, we characterised the HDX fingerprint for **6** that occupies both the αD pocket and the ATP site. We observed a similar increase in uptake in the αD loop as with **4** (Fig. 4). Compounds **4** and **6** also cause decreased uptake in the P-loop, with **6** showing more protection than **4**, similar to the protection we observed with **3**.

All these fingerprints are consistent with the known binding modalities of these ligands to CK2α and can be used to identify the binding modes of new compounds. We performed the same experiments to characterise the binding of compound **1** to CK2α (Fig. 4). Addition of **1** to CK2α causes decreased uptake in the P-loop with a pattern and magnitude of change similar to the addition of AMPPNP-Mg^2+^. However, no changes are observed in the αD pocket, as was observed with the addition of **3**. These results are consistent with the binding mode seen in the crystal structures where **1** makes significant hydrogen bonding just to Lys68 with only weak water-mediated interaction to the hinge region. These data do not allow us to unequivocally eliminate binding at the proposed allosteric site. However, if the allosteric binding mode were correct, two clear differences would be expected. Firstly, a change in the uptake by residues 67-88, part of Site A, would be expected due to the main interaction being with the acidic group of **1** (SI Fig. 8). We do not observe changes in uptake in this region despite it being redundantly covered by eight peptides (SI Fig. 11). Secondly, no change in the uptake by the P-loop would be expected since the modelling does not predict significant interactions with that part of the protein. Interactions with the P-loop, such as that seen for **1**, are one of the major features of ATP site binders, consistent with our structural characterization of **1** and **CAM4066**.

### Ligand-observed NMR analyses

To further explore the binding of the new compounds to CK2α, we have used various ligand-based NMR methods, similar to the experiments in the original characterization of these molecules. We were particularly concerned about the evidence for the simultaneous binding of **CX4945** and **1** due to the presence of strong signals for **CX4945** in the STD NMR spectra. In order to observe a signal in an STD spectrum, the saturation of protein resonances has to be transferred to the bound ligand, which then dissociates to join the pool of bulk ligand in solution before it can be detected. This means that the size of the signal for a given ligand is proportional to the amount of the ligand that binds to *and* is released by the protein during the saturation step of the STD experiment. Therefore, ligands with relatively fast association (*k*_*on*_) and dissociation (*k*_off_) rates give the largest signal.^67,68^

We have determined the off rate of **CX4945**, that binds to CK2α with low nanomolar to picomolar affinity^47^, from CK2α to be 0.0039 s^−1^ by SPR (SI Fig. 12). When ligands have *k*_off_ rates <0.1 s^−1^, the saturation cannot be transferred effectively to bulk ligand in solution resulting in no observable STD effect^68^. The experimentally determined off-rate for **CX4945** means that less than 1% of **CX4945** would have dissociated after the 2-second saturation time used in the NMR pulse sequence. This would mean that the concentration of saturated ligand in solution would be less than 0.1 μM. This difference would not be observable by NMR.

To resolve these issues, we have repeated the STD NMR studies reported in the original publication ^35^ and used two additional ligand-based NMR experiments, CPMG and Water-LOGSY^67^, for further validation. Our STD experiment shows that **1** binds to CK2α, as revealed by detectable transferred ^1^H intensities (Fig. 5A(i)). However, when the same experiment is run in the presence of **CX4945**, a significant reduction of the STD signals from **1** are observed, indicating direct competition between the two ligands (Fig. 5A(ii)). In the original paper, strong **CX4945**-derived signals are also clearly visible. The signals - which are comparable in magnitude to the much weaker binding ligand **1** - could instead arise from non-specific binding of **CX4945** to lower affinity sites on CK2α (as was shown by native mass spectrometry^35^), precipitation of **CX4945** leading to false positive signals in the experiment or binding of CX4945 to soluble aggregates of **1**.

Given the disparity in the STD NMR experiments between our results and those in the original report, we have also used CPMG and Water-LOGSY experiments. These exploit different NMR phenomena (relaxation rates, transfer of saturation) to observe the binding of the ligand to the protein^67^. Using three different methods to observe the binding of the ligand reduces the chance of experimental anomalies interfering with the outcome of the experiment. The Water-LOGSY experiment shows clear binding of **1** to CK2α and complete inhibition of binding in the presence of **CX4945** (Fig. 5B). Similarly, in the CPMG experiment, an increase in the signal from the aromatic protons of **1** was observed in the presence of **CX4945** indicating a reduction in binding of **1** to CK2α (Fig. 5C). In conclusion, all three ligand-observe NMR experiments show binding of **1** to CK2α and in all three experiments binding is abolished by **CX4945**, suggesting direct competition between these two ligands as they bind to the same site.

To further confirm the validity of our results, we repeated the CPMG NMR study using the lower affinity ligands **3, 5** and **CAM4066** as competitors (SI Fig. 13). **3** is a small, low affinity ligand in the ATP site and as such it is less likely to have an allosteric affect that might prevent binding at the adjacent Site A – an argument that could explain inhibition of binding to **1** in the presence of **CX4945**. Therefore, the reduction in the binding of **1** in the presence of **3** (SI Fig. 13B) is a more convincing proof that **1** is binding in the ATP site. **CAM4066** links the ATP site and the αD site and, as predicted, reduces the binding of **1** to CK2α (SI Fig. 13C). To rule out binding of **1** to the αD site, we have also used **5**, an optimized fragment with high affinity for the αD site. This should not prevent binding in the ATP site. Indeed, ligands similar to **5** have been shown to bind simultaneously with ATP site fragments by X-ray crystallography (PDB:6EHK)^26^. As predicted, and unlike with **CX4945, 3** and **CAM4066**, the αD site ligand **5** has no effect on the binding of **1** to CK2α (SI Fig. 13D). Similar CPMG and STD NMR experiments were performed with the CK2α KKK/AAA mutant with the same outcomes (SI Fig. 14 and 15). These results further demonstrate that **1** binds in the ATP site.

### Isothermal titration calorimetry

As a final confirmation, we have used isothermal titration calorimetry to determine affinities of **1** and **2** to CK2α and to evaluate the effect of competing ligands and CK2α mutations. This process has been successfully used to verify the binding site of novel CK2 inhibitors that bind in the αD site^26^. The affinities of compounds **1** and **2** for CK2α were determined to be 22 μM and 2.2 μM respectively (Fig. 6A and C). When the same experiments were performed in the presence of saturating concentrations of **CX4945**, no binding of either compound to CK2α was observed (Fig. 6B and 6D). This observation is in line with all our previous data and suggests that compounds **1** and **2** bind in the same site as **CX4945**. We also performed the experiment with **2** in the presence of **CAM4066**, which competes with ATP, but uses only a small benzoic acid moiety to interact with the ATP site, deriving most of its binding energy from the αD pocket. **CAM4066** abolished the binding of **2** to CK2α (SI Fig. 16). In the modelling of the binding of **1** and **2** reported previously^35,36^, Lys74 was proposed to form a vital part of the binding site and to interact with the carboxylic acid of **1** and **2**. Therefore, mutations to this residue should reduce the affinity of **1** and **2** for CK2α. To test this, we repeated the ITC experiments using the construct CK2α KKK/AAA, in which three adjacent lysine residues (Lys74-76) have been mutated to alanine. These mutations had a negligible effect on the affinity of **1** or **2** for CK2α as we determined near identical affinities of 27 μM and 3.3 μM, respectively (SI Fig. 16). The binding of **1** and **2** to CK2α KKK/AAA was also fully inhibited by the presence of **CX4945** and **CAM4066** respectively (SI Fig. 16). These results are further evidence that the ATP site, not the proposed allosteric site, is the primary binding site for **1** and **2**.

**Figure 6.**
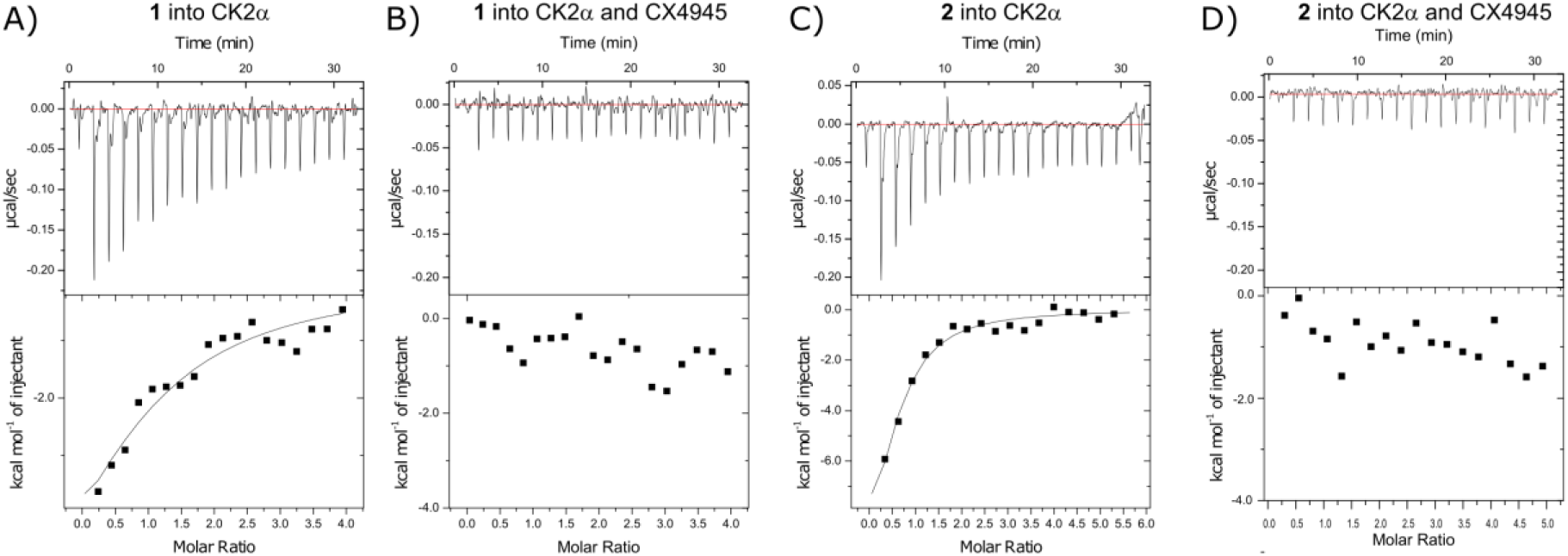
ITC measurement of binding of 1 and 2 to CK2α. (A) Titration of 1 into 10 μM CK2α Titration of 2 into 10 μM CK2α. (B) Titration of **1** into 10 μM CK2α Titration of **2** into 10 μM CK2α in the presence of **CX4945** (C) Titration of **2** into 20 μM CK2α. (D) Titration of **2** into 20 μM CK2α in the presence of 100 μM **CX4945**.

## Discussion

We present compelling and consistent evidence that **1** and **2** bind in the ATP site of CK2α, not in Site A, as was previously proposed.^35,36^ We see no evidence for binding in Site A. This is as expected, based on our analysis of the structural data of similar kinase inhibitors. In the original study of Series A,^36^ five main pieces of evidence were used to verify a novel binding mode. This evidence included: competitive native mass spectroscopy studies, ligand-based NMR, thermal shift analysis and biochemical assays. While we were preparing this manuscript for publication, the Niefind group published their analysis of the same series of compounds, which also showed binding to the ATP site by X-ray crystallography.^69^ While they have not done other biophysical analyses on the binding, they do show by enzymatic assay that these inhibitors are ATP competitive, consistent with our data.

One of the cornerstones of the theory of allosteric inhibition was the observation that that **CX4945** binds CK2α simultaneously with **1**, and that **1** competes with inositol hexaphosphate (IP_6_, phytic acid), shown by mass spectrometry. The data presented indicate that at least three molecules of **1** bind to CK2α. In the presence of **CX4945** this is reduced to two binding sites, but no data were presented to confirm the identity of these sites in which **1** was binding when it bound simultaneously with **CX4945** (for example mutational studies combined with native mass spectrometry). Some of these sites are expected to be non-specific, low affinity sites and the mass spectrometric data does not confirm which is the primary binding site of **1**. The first evidence presented for the location of the binding site of these compounds is the observation that in the presence of **1** the complex between IP_6_ and CK2α is not observed in the native mass spectra. This led to the theory that **1** binds in the proposed site as a crystal structure was published by Lee et al (PDB:3W8L)^70^ of IP_6_ apparently binding to CK2α in this site. However, the binding site of IP_6_ presented in that crystal structure is more controversial than is discussed in the publication.^70^ In this structure, IP_6_ is found at a crystal contact where half of the site is composed of residues from a symmetry related CK2α molecule in the crystal lattice (SI Fig. 17). Therefore, IP_6_ could in theory bind to both these sites. From this crystal structure the binding site could be located around Lys74 or His276. However, as the site is only fully formed in a crystal lattice (SI Fig. 17), IP_6_ may not in actual fact bind to CK2α at either of these sites in solution. IP_6_ is also seen to bind weakly at a site centered on His236 in several other crystal structures (PDB:5OQU, 5ORJ, 5MPJ, 5CVH)^26,27,60^. The native mass spectrometric competition studies with **CX4945** or IP_6_ do not therefore unambiguously confirm the binding mode of **1**.

Another cornerstone of the allosteric site binding was the observation of the binding by ligand-based NMR and simultaneous detection of **CX4945** binding, interpreted as lack of competition. As discussed before, a high affinity ligand like **CX4945**, with slow dissociation rate should not yield detectable signal in these experiments. The binding of **CX4945** observed in those STD experiments is however more likely to be due to an artefact in the experimental setup. This could be through a number of mechanisms including aggregation of the ligand or binding to non-specific lower affinity binding sites outside the ATP site.^71^ These lower affinity sites would have higher on/off rates than the ATP site and therefore show a larger signal in the STD spectrum. Non-specific and low affinity sites can have a confusing influence on competition studies. This is a particular problem with the NMR studies as ligand-observed NMR is optimized for studying low affinity sites and requires ligands with rapid on/off rates in order for the signal (and therefore binding) to be observed. Therefore, blocking the primary (high affinity) site may not remove the binding signal arising from other sites.

The competition studies in the original papers were done at 500 μM ligand concentration, well beyond the observed affinities or K_i_s of **1** and **2** and in a range where non-specific binding would be more likely to occur.^35,36^ The binding signals observed for **CX4945** in the STD spectra cast some doubt on the conclusions drawn about the simultaneous binding of **1** and **CX4945**. This is because the signals observed for **CX4945** cannot be due to binding in the ATP site as the dissociation rate for this site is too slow. The major difference between our experimental system and the system reported in the literature is the DMSO content. In brief, in the procedure reported here stocks of **1** and **2** were made in d_6_-DMSO and diluted 20-fold in the buffer. This was done to ensure the ligands were fully soluble in the experiment and led to a final d_6_-DMSO content of 5% (which was also used to provide the field-frequency lock signal). When we first attempted to run the experiment without DMSO, we observed precipitation of the ligands. As aggregation can lead to false positives in STD experiments it was decided to use DMSO in all our experiments.^71^ However, in the original experiment D_2_O was used for the lock and no DMSO was used to solubilise the ligand. The difference in the solvent conditions may mean that **1** and **2** were closer to the limits of their solubility in the original experiments and thus more likely to form aggregates. Aggregates are common causes of false positives in biochemical assays and are often missed as they may not be easily visible to the naked eye. These aggregates may lead to the signals observed in the STD for **CX4945** and **1**.

The biochemical assays performed previously showed that the compounds exhibited mixed mode non-competitive behavior and led to the understandable conclusion that they do not bind in the ATP site. However, this is contradictory to the binding mode we see in the crystal structures. A proposed explanation for the non-competitive inhibition observed in the assay comes from the thermal stability data also presented in the original paper.^35^ The data show that the thermal stability of the proteins decreases in the presence of **1** and **2**. This thermal de-stabilization could be due to a number of different mechanisms caused by specific and non-specific effects. These mechanisms include; breaking up of protein complexes by specific binding to the protein; binding to the unfolded or partially unfolded protein; the formation of aggregates that bind the protein and unfold it and specific binding that traps the protein in a less stable conformation^72–75^.

It is impossible that the first mechanism is operating in this example as CK2α is present as a monomer. The other mechanisms however may explain the conflicting results presented. All proteins exist in an equilibrium between the native and unfolded form of the protein.^76^ For the majority of ligands, the ligand binds to the native form of the protein which stabilizes the folded form leading to a positive thermal shift. If the ligand binds to the unfolded or partially unfolded form of the protein and stabilize the unfolded form this could lead to a destabilization of the protein. This would in turn lead to a decrease in the *T*_m_. Binding to the unfolded protein can occur by two mechanisms, the first mechanism is by the formation of colloidal aggregates of the small molecules, onto which the protein would then be sequestered.^75^ The aggregates are hydrophobic and preferentially bind the unfolded or partially unfolded form of the protein where the hydrophobic core is exposed. Therefore, a negative shift in the *T*_m_ is observed. This is one of the most common forms of non-specific inhibition that plague biochemical assays and would be seen in assays as noncompetitive inhibition. The second mechanism occurs when the ligand monomers bind to a partially unfolded or fully unfolded form of the protein stabilizing that form. It may be possible for the binding to the partially unfolded form to be a specific form of inhibition.

Modelling experiments proposed Site A as the binding site for **1** and **2** (SI Fig. 8)^35^. We have accumulated extensive evidence that these compounds bind in the ATP site. However, Site A as a general ligand binding site has been verified by our structures of ADP and GDP bound in this location. This suggests that Site A could be utilized for the development of novel inhibitors of CK2α and it also demonstrates that this site is accessible for ligand binding in our crystal forms. If this site were to be successfully targeted by inhibitors, it would require extensive structural and biochemical validation to confirm that compounds are really binding in the new site and not in one of the several known small molecule binding sites in CK2α.

We believe that the data we have presented here leads to the conclusion that the compounds represented by **1** and **2** bind to the ATP site and will inhibit CK2α via an orthosteric, ATP-competitive mechanism. This diverse set of data pointing to **1** and **2** binding at the ATP site includes multiple crystal structures, competitive ITC studies, competitive NMR studies, H/D exchange mass spectrometry and computational analyses of the ligand structures. We also believe that the evidence presented previously in support of an allosteric mode of inhibition for these compounds may have been distorted by non-specific binding of the compounds to CK2α, which in turn affected the interpretation of competition data with **CX4945**.

## Supporting information

HDX dataset

## Acknowledgements

We would like to thank Dr Glyn Williams and Teodors Pantelejevs for their critical comments on the manuscript. We are grateful for Diamond Light Source for access to beamlines i04, i04-1 and i24 (proposals: mx9537, mx14043, mx18548). We thank the X-ray Crystallographic and Biophysics Facilities for access to instrumentation. We thank Apollo Therapeutics that funds our CK2α inhibitor development project for allowing the use of the data on CX4945 binding kinetics for this publication. The work in the D’Arcy group was funded by the NIH National Institute of General Medical Sciences (GM133751) and start-up funds to S. D’Arcy.

## Structural coordinates in Protein Data Bank

6YPG (CK2α in complex with **2**), 6YPH (CK2α in complex with **2**), 6YPJ (CK2α in complex with **1**), 6YPK (CK2α in complex with **GDP**), 6YPN (CK2α in complex with **ATP**).

