## Supplemental Information for "Proposed allosteric inhibitors bind to the ATP site of CK2α"

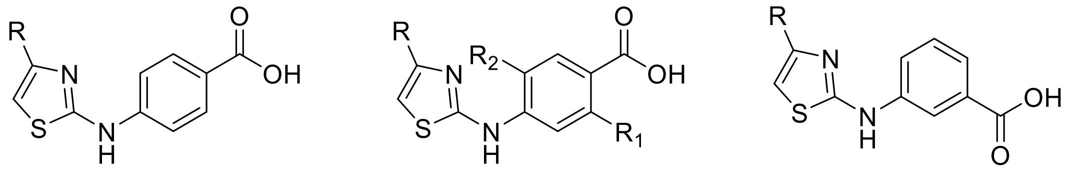

**Supplementary Figure 1.** The series of proposed allosteric inhibitors of CK2α developed by Bestegen et al. (Series A)[1].

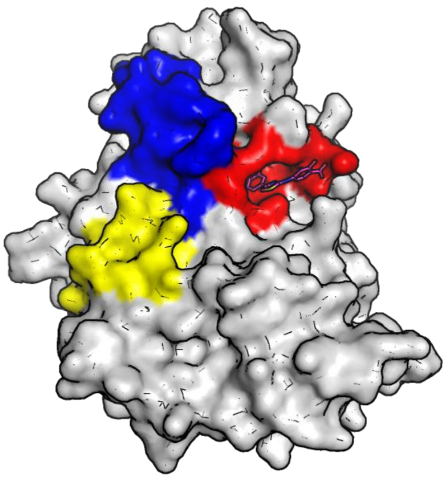

**Supplementary Figure 2.** Position of the proposed allosteric site, referred to as Site A (highlighted in red), based on the modelling of the complex of **1** with CK2α in the original publications[1]. The location of the ATP site (blue) and the αD site (yellow) are also shown.

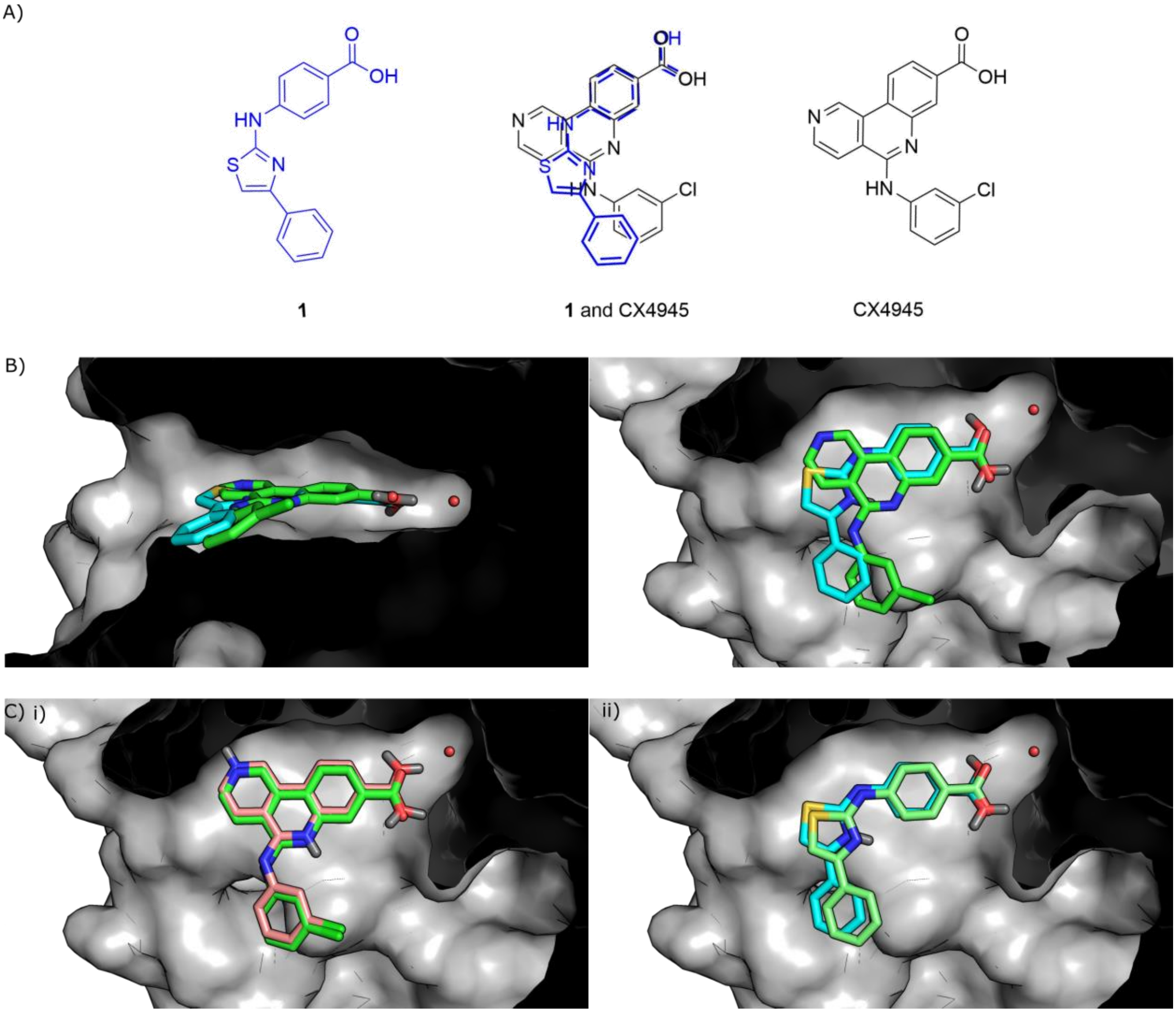


**Supplementary Figure 2**: Comparison of **1** and **CX4945** by overlay and modelling A) Overlay of the structures of **1** and **CX4945**. B)The binding mode of **1** (blue) to CK2α predicted by GOLD with the binding mode of **CX4945** (green) observed in the crystal structure (pdb:3PE1) superimposed on the model[2]. C) i)The binding mode of **CX4945** to CK2α predicted by GOLD with the binding mode of **CX4945** determined in the crystal structure (pdb:3PE1)[2] superimposed on the model. The model clearly predicts the correct binding mode ii) The binding mode of **1** to CK2α predicted by GOLD with the binding mode of **1** determined in the crystal structure (pdb:6YPJ) superimposed on the model. The modelling clearly predicts the correct binding mode.

**Supplementary Table 1.** Ligand structures and SMILES strings from PDB search with search terms: "E.C. number 2.7.11.1 +benzoic"

| PDB | Structure | Binding site |
|---|---|---|
| 2E9O | 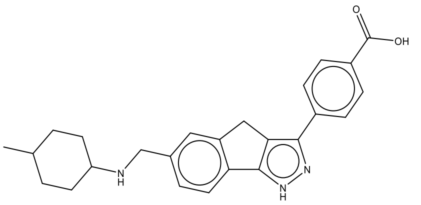<br>CC1CCC(CC1)NCc2ccc-3c(c2)Cc4c3[nH]nc4c5ccc(cc5)C(=O)O | ATP |
| 2WQO | 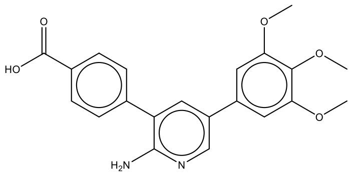<br>COc1cc(cc(c1OC)OC)c2cc(c(nc2)N)c3ccc(cc3)C(=O)O | ATP |
| 2WTV | 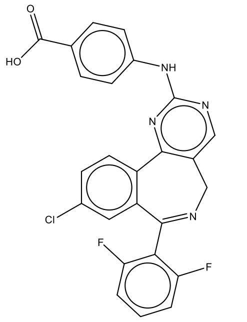<br>c1cc(c(c(c1)F)C2=NCc3cnc(nc3-c4c2cc(cc4)Cl)Nc5ccc(cc5)C(=O)O)F | ATP |

| | | |
|---|---|---|
| 2X7G | 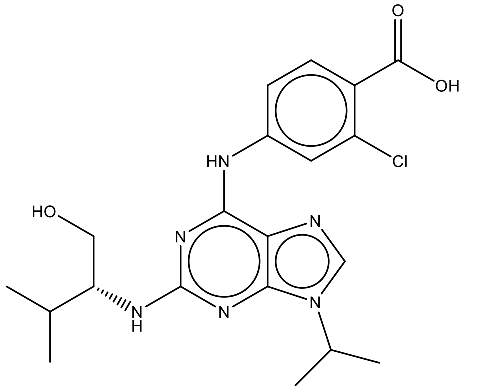 CC(C)[C@H](CO)Nc1nc(c2c(n1)n(cn2)C(C)C)Nc3ccc(c(c3)Cl)C(=O)O | ATP |
| 2XK4 | 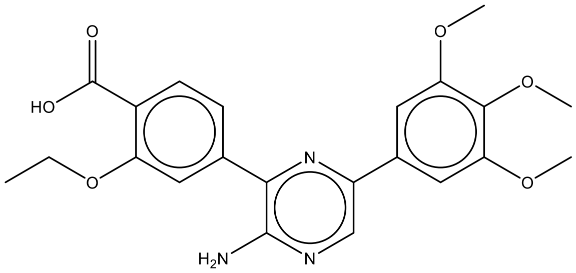 CCOc1cc(ccc1C(=O)O)c2c(ncc(n2)c3cc(c(c(c3)OC)OC)OC)N | ATP |
| 2XK8 | 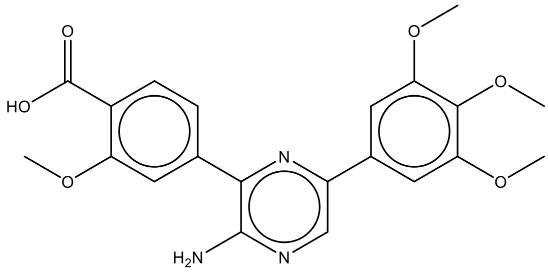 COc1cc(ccc1C(=O)O)c2c(ncc(n2)c3cc(c(c(c3)OC)OC)OC)N | ATP |
| 2XKC | 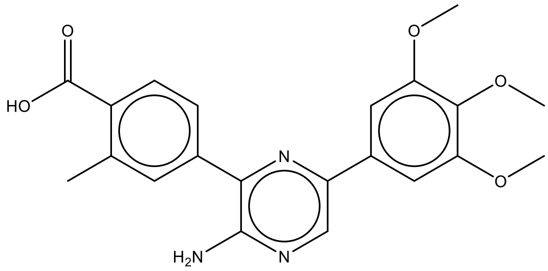 Cc1cc(ccc1C(=O)O)c2c(ncc(n2)c3cc(c(c(c3)OC)OC)OC)N | ATP |

| | | |
|---|---|---|
| 2XKD | 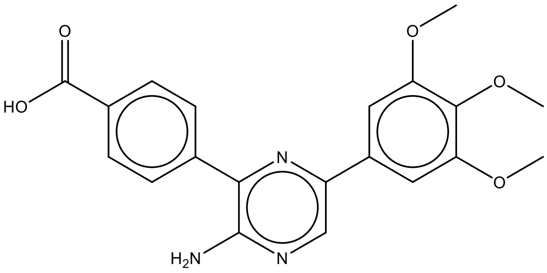<br>COc1cc(cc(c1OC)OC)c2cnc(c(n2)c3ccc(cc3)C(=O)O)N | ATP |
| 3AXW | 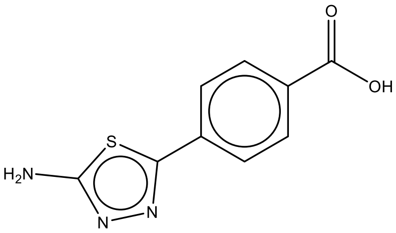<br>c1cc(ccc1c2nnc(s2)N)C(=O)O | ATP |
| 3BGZ | 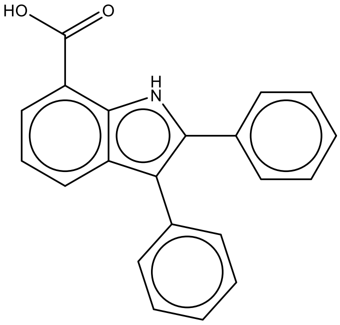<br>c1ccc(cc1)c2c3cccc(c3[nH]c2c4ccccc4)C(=O)O | ATP |
| 3BQR | 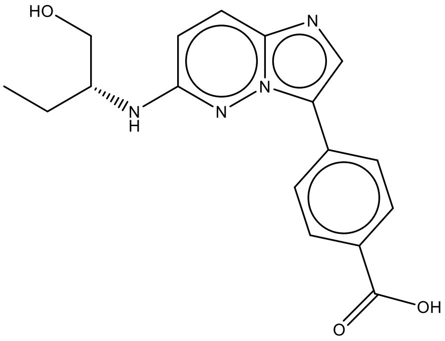<br>CC[C@H](CO)Nc1ccc2ncc(n2n1)c3ccc(cc3)C(=O)O | ATP |

| | | |
|---|---|---|
| 3HDM | 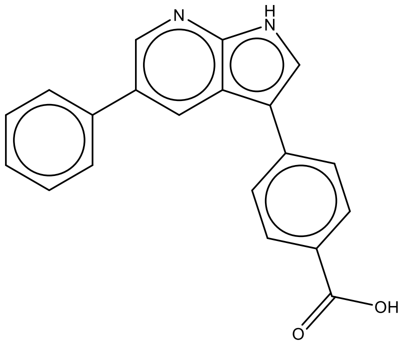 c1ccc(cc1)c2cc3c(c[nH]c3nc2)c4ccc(cc4)C(=O)O | ATP |
| 3NGA (CX4945) | 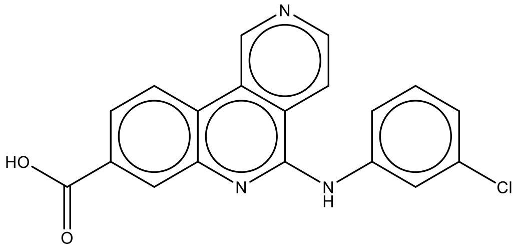 c1cc(cc(c1)Cl)Nc2c3ccncc3c4ccc(cc4n2)C(=O)O | ATP |
| 3PE2 | 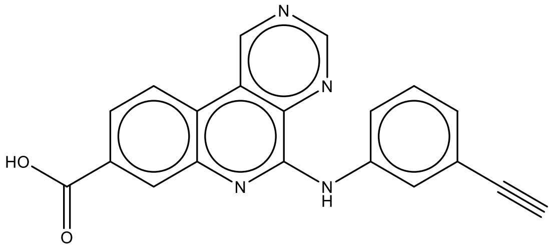 C#Cc1cccc(c1)Nc2c3c(cncn3)c4ccc(cc4n2)C(=O)O | ATP |
| 3R0T | 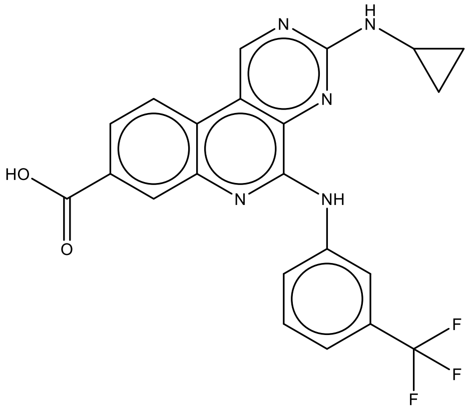 c1cc(cc(c1)Nc2c3c(cnc(n3)NC4CC4)c5ccc(cc5n2)C(=O)O)C(F)(F)F | ATP |
| 3UNZ | 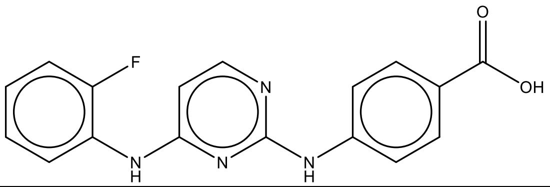 | ATP |

| | | | |
|---|---|---|---|
| | c1ccc(c(c1)Nc2ccnc(n2)Nc3ccc(cc3)C(=O)O)F | | |
| 3UO4 | 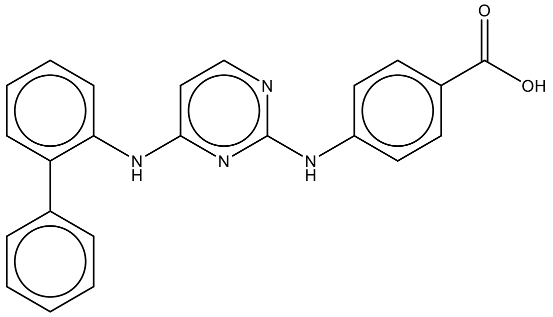<br><br>c1ccc(cc1)c2ccccc2Nc3ccnc(n3)Nc4ccc(cc4)C(=O)O | | ATP |
| 3UO5 | 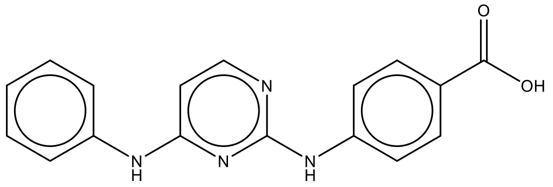<br><br>c1ccc(cc1)Nc2ccnc(n2)Nc3ccc(cc3)C(=O)O | | ATP |
| 3UO6 | 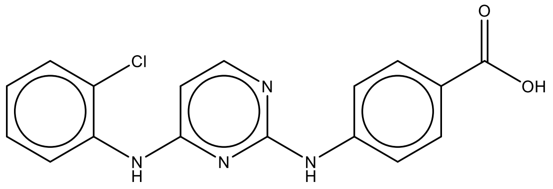<br><br>c1ccc(c(c1)Nc2ccnc(n2)Nc3ccc(cc3)C(=O)O)Cl | | ATP |
| 3UOD | 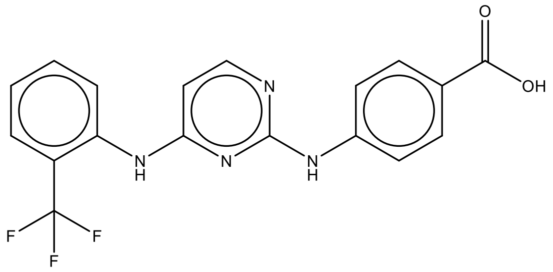<br><br>c1ccc(c(c1)C(F)(F)F)Nc2ccnc(n2)Nc3ccc(cc3)C(=O)O | | ATP |
| 3UOH | 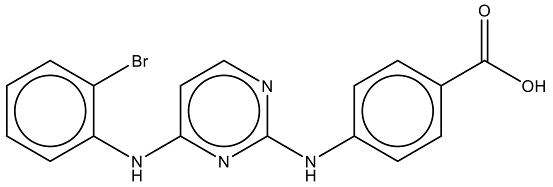<br><br>c1ccc(c(c1)Nc2ccnc(n2)Nc3ccc(cc3)C(=O)O)Br | | ATP |

| | | |
|---|---|---|
| 3UOJ | 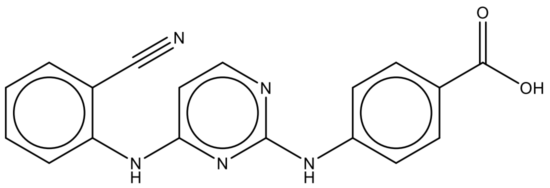 c1ccc(c(c1)C#N)Nc2ccnc(n2)Nc3ccc(cc3)C(=O)O | ATP |
| 3UOK | 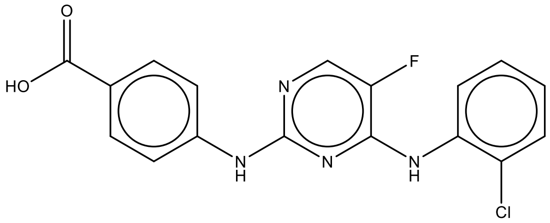 c1ccc(c(c1)Nc2c(cnc(n2)Nc3ccc(cc3)C(=O)O)F)Cl | ATP |
| 3UP2 | 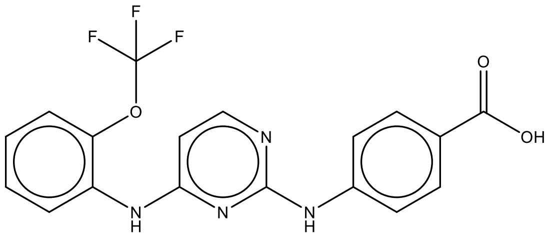 c1ccc(c(c1)Nc2ccnc(n2)Nc3ccc(cc3)C(=O)O)OC(F)(F)F | ATP |
| 3UP7 | 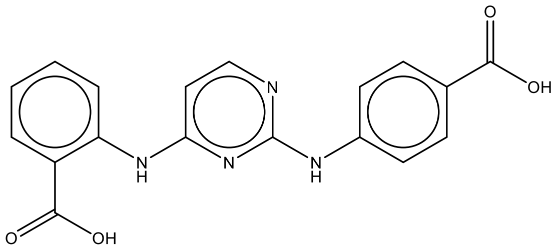 c1ccc(c(c1)C(=O)O)Nc2ccnc(n2)Nc3ccc(cc3)C(=O)O | ATP |
| 4AW2 | 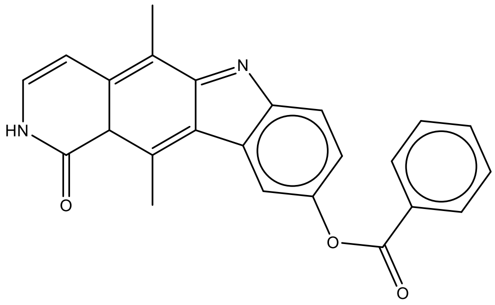 CC1=C2c3cc(ccc3N=C2C(=C4C1C(=O)NC=C4)C)OC(=O)c5ccccc5 | Interface site |

| | | |
|---|---|---|
| 4BTM | 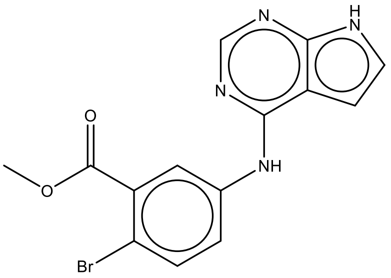<br>COC(=O)c1cc(ccc1Br)Nc2c3cc[nH]c3ncn2 | ATP |
| 4CFE | 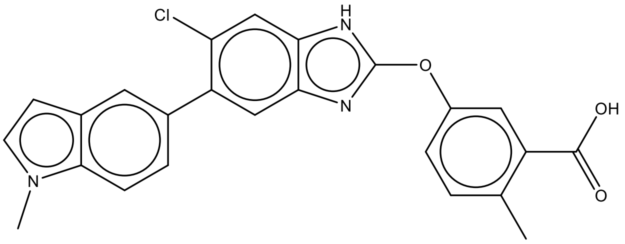<br>Cc1ccc(cc1C(=O)O)Oc2[nH]c3cc(c(cc3n2)c4ccc5c(c4)ccn5C)Cl | Interface site |
| 4DEA | 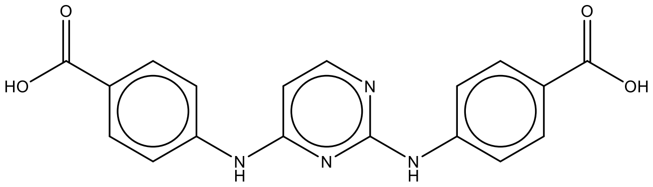<br>c1cc(ccc1C(=O)O)Nc2ccnc(n2)Nc3ccc(cc3)C(=O)O | ATP |
| 4FT0 | 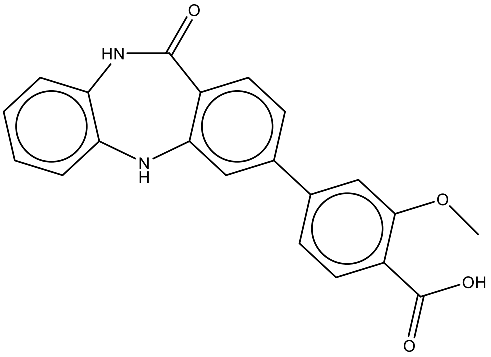<br>COc1cc(ccc1C(=O)O)c2ccc3c(c2)Nc4ccccc4NC3=O | ATP |
| 4FTA | 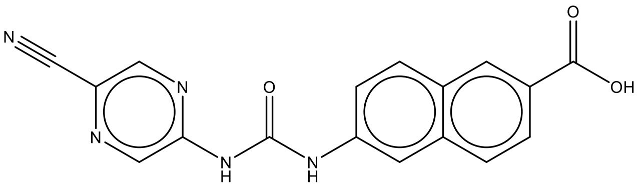<br>c1cc(cc2c1cc(cc2)NC(=O)Nc3cnc(cn3)C#N)C(=O)O | ATP |

| | | |
|---|---|---|
| 4QYE | 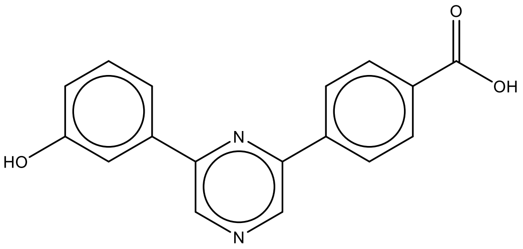<br>c1cc(cc(c1)O)c2cncc(n2)c3ccc(cc3)C(=O)O | ATP |
| 4QYF | 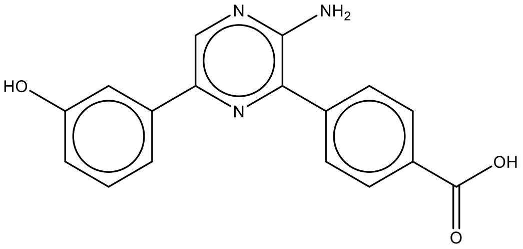<br>c1cc(cc(c1)O)c2cnc(c(n2)c3ccc(cc3)C(=O)O)N | ATP |
| 4RLK | 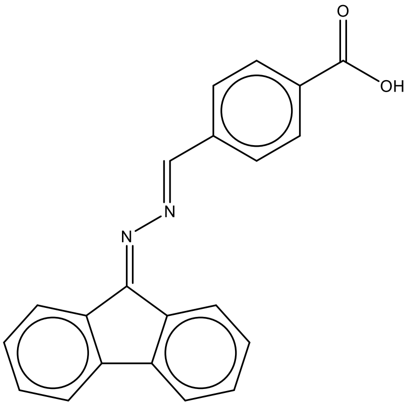<br>c1ccc2c(c1)-c3ccccc3C2=N/N=C/c4ccc(cc4)C(=O)O | ATP |
| 4UB7 | 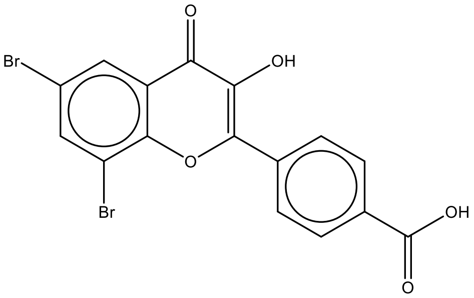<br>c1cc(ccc1C2=C(C(=O)c3cc(cc(c3O2)Br)Br)O)C(=O)O | ATP |
| 4WOT | 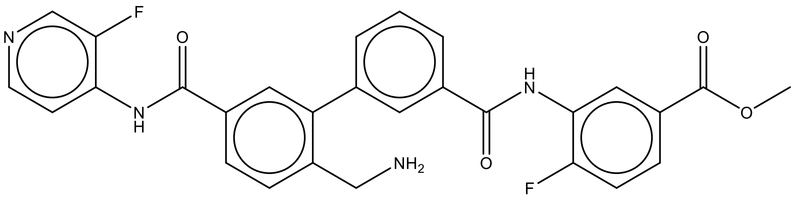<br>COC(=O)c1ccc(c(c1)NC(=O)c2cccc(c2)c3cc(ccc3CN)C(=O)Nc4ccncc4F)F | ATP |

| | | |
|---|---|---|
| 5B0X | 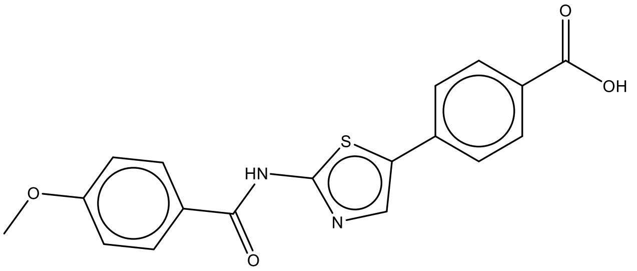 COc1ccc(cc1)C(=O)Nc2ncc(s2)c3ccc(cc3)C(=O)O | ATP |
| 5C5E | 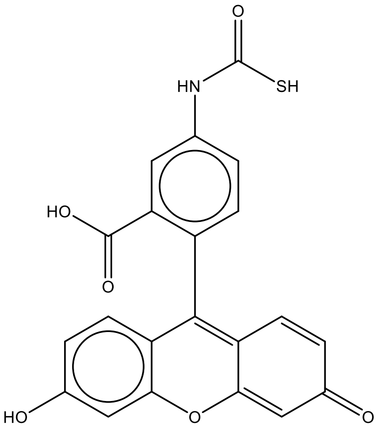 c1cc(c(cc1NC(=O)S)C(=O)O)C2=C3C=CC(=O)C=C3Oc4c2ccc(c4)O | ATP |
| 5CSP | 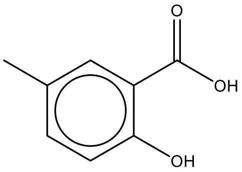 Cc1ccc(c(c1)C(=O)O)O | ATP |
| 5CSV | 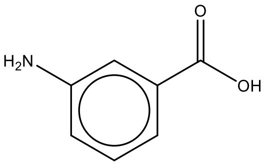 c1cc(cc(c1)N)C(=O)O | ATP |
| 5CU3 | 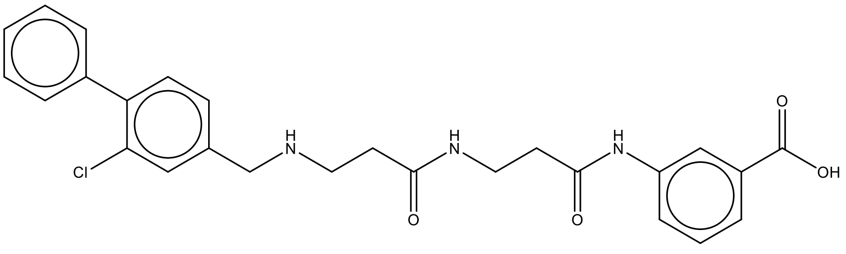 c1ccc(cc1)c2ccc(cc2Cl)CNCCC(=O)NCCC(=O)Nc3cccc(c3)C(=O)O | ATP |

| | | |
|---|---|---|
| 5M4F | 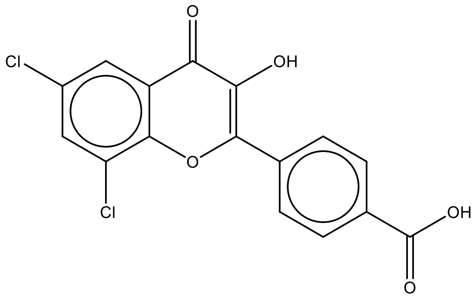<br>c1cc(ccc1C2=C(C(=O)c3cc(cc(c3O2)Cl)Cl)O)C(=O)O | ATP |
| 5MAF | 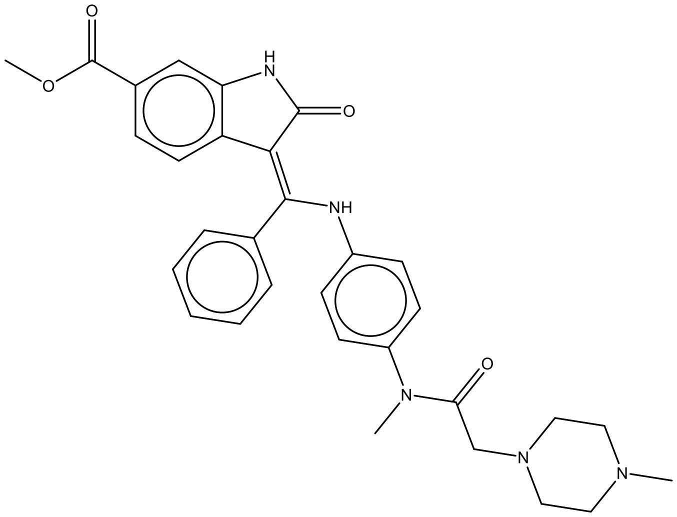<br>CN1CCN(CC1)CC(=O)N(C)c2ccc(cc2)N/C(=C\3/c4ccc(cc4NC3=O)C(=O)OC)/c5ccccc5 | ATP |
| 5MO8 | 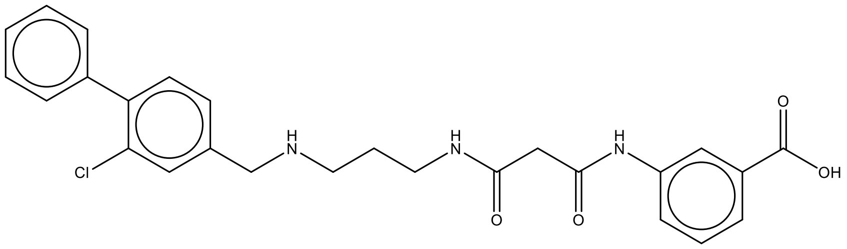<br>c1ccc(cc1)c2ccc(cc2Cl)CNCCCNC(=O)CC(=O)Nc3cccc(c3)C(=O)O | ATP |
| 5OUE | 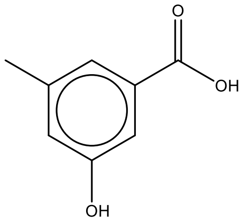<br>Cc1cc(cc(c1)O)C(=O)O | ATP |

| | | |
|---|---|---|
| 6B1U | 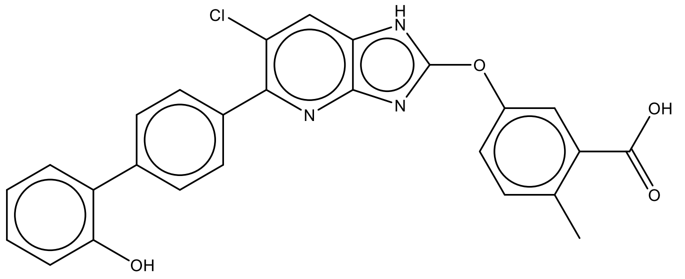<br>Cc1ccc(cc1C(=O)O)Oc2[nH]c3cc(c(nc3n2)c4ccc(cc4)c5ccccc5O)Cl | Interface site |
| 6BSK | 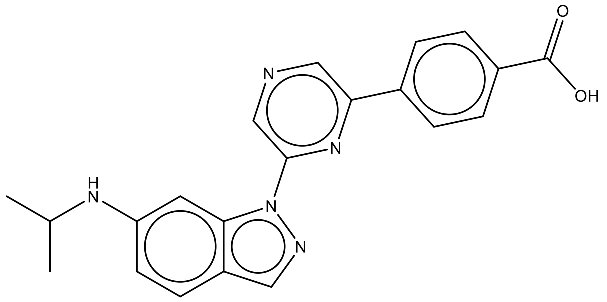<br>CC(C)Nc1ccc2cnn(c2c1)c3cncc(n3)c4ccc(cc4)C(=O)O | ATP |
| 6Q38 | 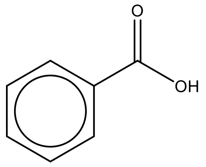<br>c1ccc(cc1)C(=O)O | ATP |

**Supplementary Table 2.** Ligand structures and SMILES strings from PDB search with search terms: "2.7.11.1 + amino benzoic acid".

| PDB | Structure | Binding site |
|---|---|---|
| 2WTV | 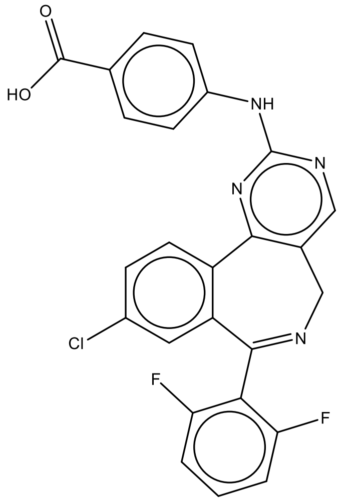<br>c1cc(c(c(c1)F)C2=NCc3cnc(nc3-c4c2cc(cc4)Cl)Nc5ccc(cc5)C(=O)O)F | ATP |
| 2X7G | 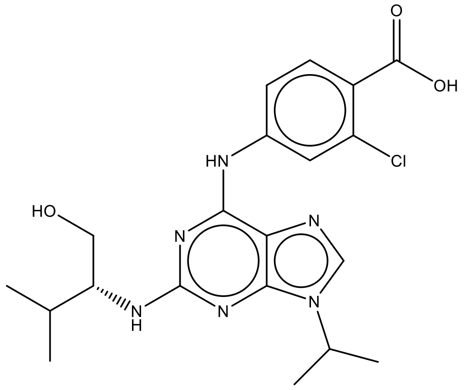<br>CC(C)[C@H](CO)Nc1nc(c2c(n1)n(cn2)C(C)C)Nc3ccc(c(c3)Cl)C(=O)O | ATP |
| 3UNZ | 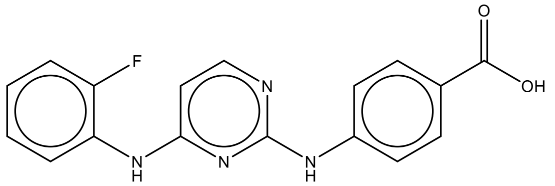<br>c1ccc(c(c1)Nc2ccnc(n2)Nc3ccc(cc3)C(=O)O)F | ATP |
| 3UO4 | 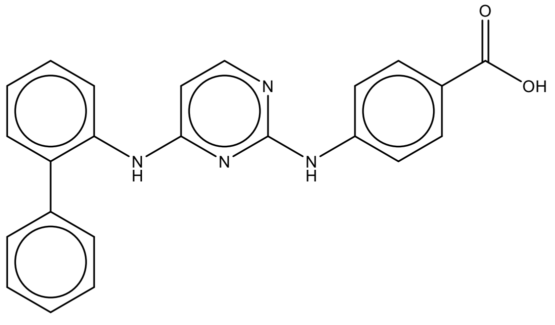<br>c1ccc(cc1)c2ccccc2Nc3ccnc(n3)Nc4ccc(cc4)C(=O)O | ATP |

| | | |
|---|---|---|
| 3UO5 | 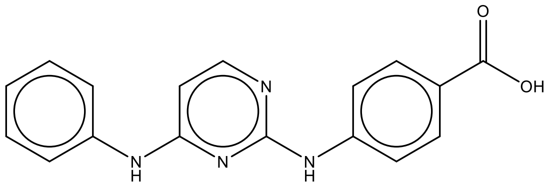<br>c1ccc(cc1)Nc2ccnc(n2)Nc3ccc(cc3)C(=O)O | ATP |
| 3UO6 | 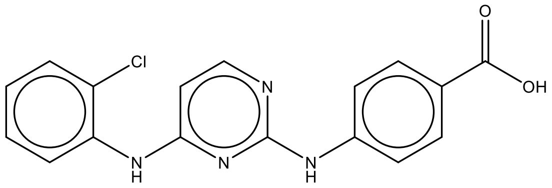<br>c1ccc(c(c1)Nc2ccnc(n2)Nc3ccc(cc3)C(=O)O)Cl | ATP |
| 3UOD | 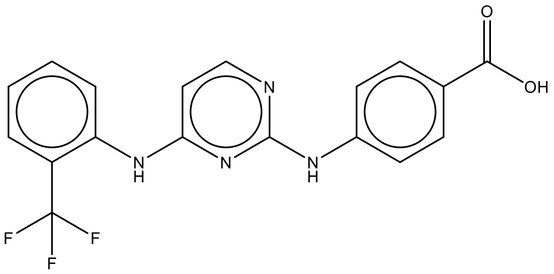<br>c1ccc(c(c1)C(F)(F)F)Nc2ccnc(n2)Nc3ccc(cc3)C(=O)O | ATP |
| 3UOH | 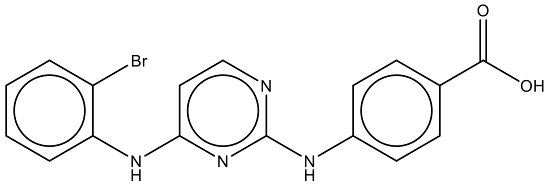<br>c1ccc(c(c1)Nc2ccnc(n2)Nc3ccc(cc3)C(=O)O)Br | ATP |
| 3UOJ | 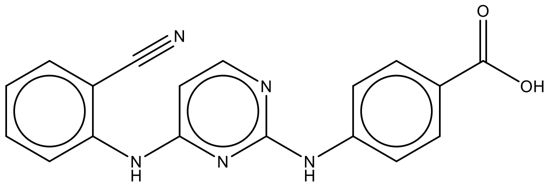<br>c1ccc(c(c1)C#N)Nc2ccnc(n2)Nc3ccc(cc3)C(=O)O | ATP |
| 3UOK | 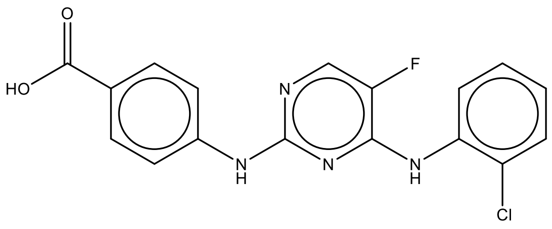<br>c1ccc(c(c1)Nc2c(cnc(n2)Nc3ccc(cc3)C(=O)O)F)Cl | ATP |

| | | |
|---|---|---|
| 3UP2 | 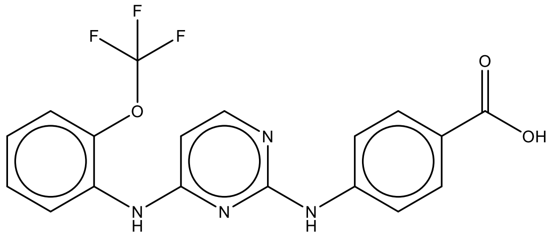<br><br>c1ccc(c(c1)Nc2ccnc(n2)Nc3ccc(cc3)C(=O)O)OC(F)(F)F | ATP |
| 3UP7 | 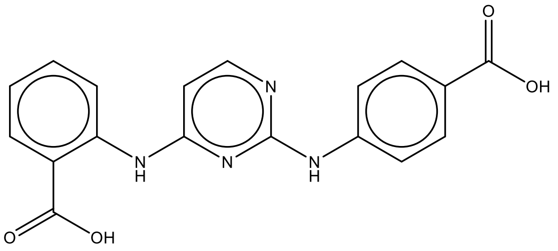<br><br>c1ccc(c(c1)C(=O)O)Nc2ccnc(n2)Nc3ccc(cc3)C(=O)O | ATP |
| 4DEA | 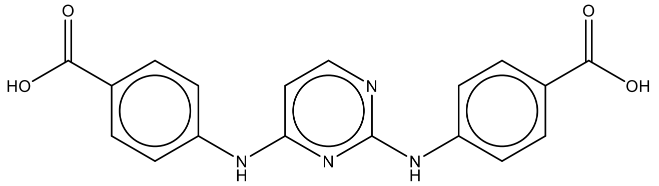<br><br>c1cc(ccc1C(=O)O)Nc2ccnc(n2)Nc3ccc(cc3)C(=O)O | ATP |

**Supplementary Table 3.** Ligand structures and SMILES strings from PDB search with search terms: “Casein Kinase II + benzoic acid”.

| PDB | Ligand structure |
|---|---|
| 3AXW | 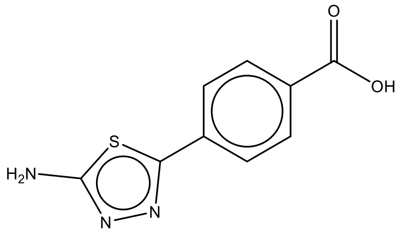<br><br>c1cc(ccc1c2nnc(s2)N)C(=O)O |
| 3NGA (CX4945 ) | 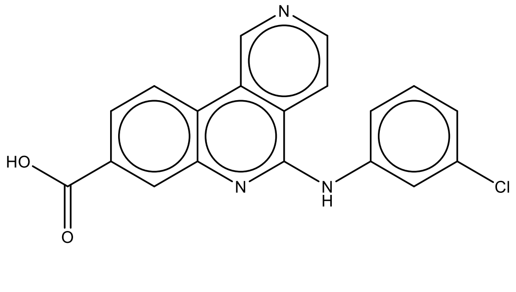<br><br>c1cc(cc(c1)Cl)Nc2c3ccncc3c4ccc(cc4n2)C(=O)O |

| | |
|---|---|
| 3PE2 | 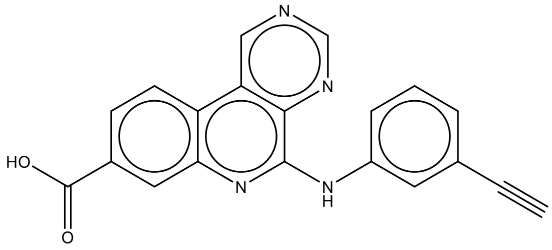<br>C#Cc1cccc(c1)Nc2c3c(cncn3)c4ccc(cc4n2)C(=O)O |
| 3R0T | 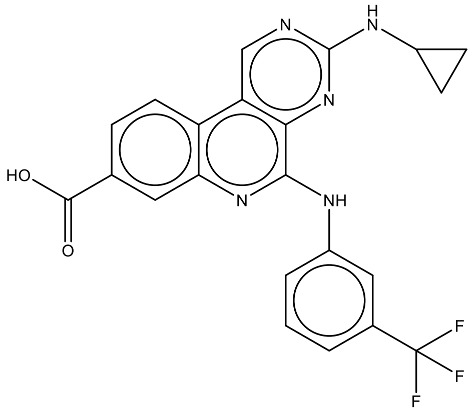<br>c1cc(cc(c1)Nc2c3c(cnc(n3)NC4CC4)c5ccc(cc5n2)C(=O)O)C(F)(F)F |
| 4RLK | 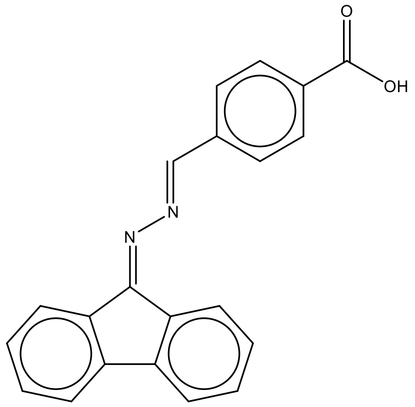<br>c1ccc2c(c1)-c3ccccc3C2=N/N=C/c4ccc(cc4)C(=O)O |
| 4UB7 | 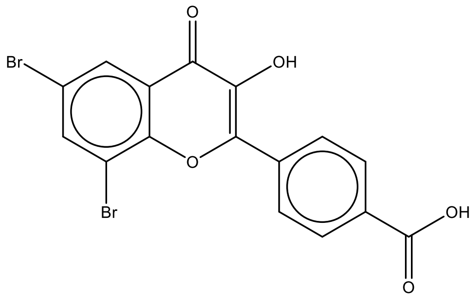<br>c1cc(ccc1C2=C(C(=O)c3cc(cc(c3O2)Br)Br)O)C(=O)O |

| | |
|---|---|
| 5B0X | 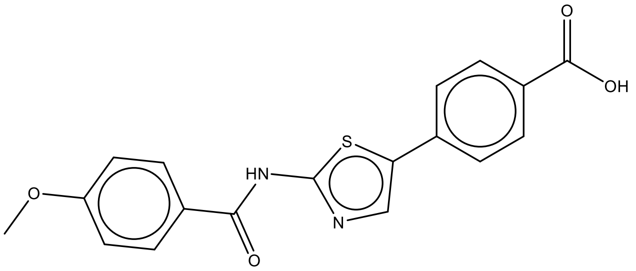 COc1ccc(cc1)C(=O)Nc2ncc(s2)c3ccc(cc3)C(=O)O |
| 5CSP | 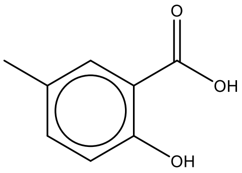 Cc1ccc(c(c1)C(=O)O)O |
| 5CSV | 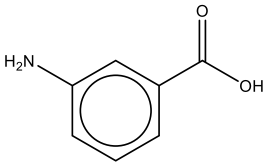 c1cc(cc(c1)N)C(=O)O |
| 5CU3 | 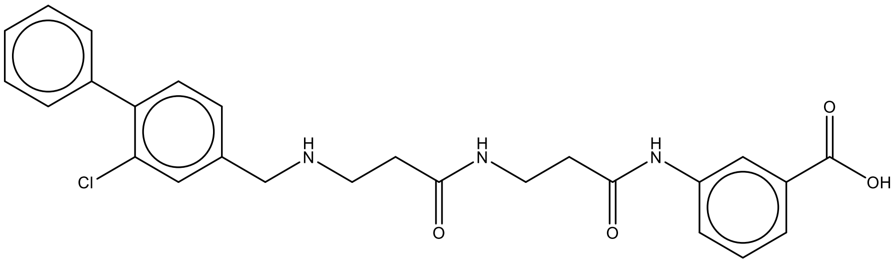 c1ccc(cc1)c2ccc(cc2Cl)CNCCC(=O)NCCC(=O)Nc3cccc(c3)C(=O)O |
| 5M4F | 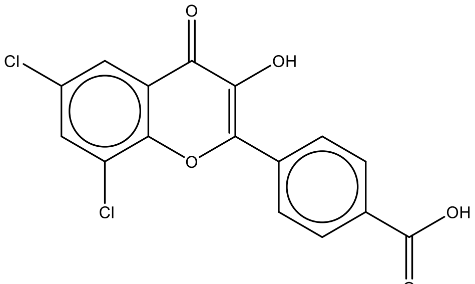 c1cc(ccc1C2=C(C(=O)c3cc(cc(c3O2)Cl)Cl)O)C(=O)O |

| | |
|---|---|
| 5MO8 | 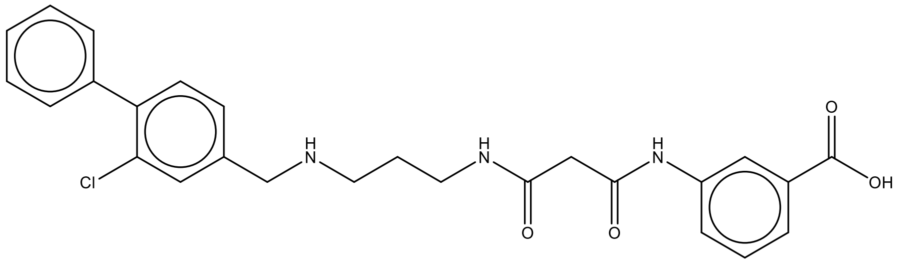<br>c1ccc(cc1)c2ccc(cc2Cl)CNCCCNC(=O)CC(=O)Nc3cccc(c3)C(=O)O |
| 5OUE | 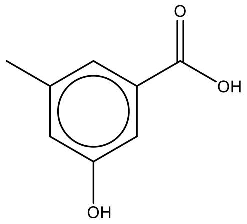<br>Cc1cc(cc(c1)O)C(=O)O |
| 6Q38 | 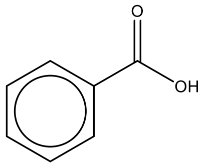<br>c1ccc(cc1)C(=O)O |

**Supplementary Table 4.** Ligand structures and SMILES strings from PDB search with terms: "Casein Kinase II + benzoic acid + hinge binder".

| PDB | Ligand structure |
|---|---|
| 3AXW | 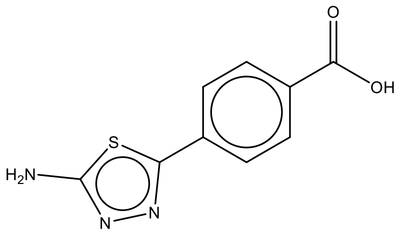<br>c1cc(ccc1c2nnc(s2)N)C(=O)O |
| 3NGA (CX4945) | 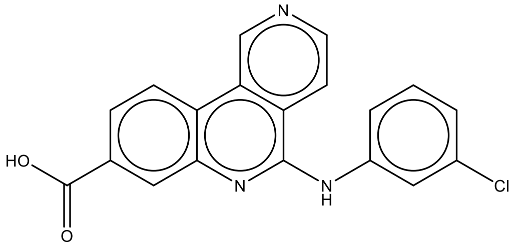<br>c1cc(cc(c1)Cl)Nc2c3ccncc3c4ccc(cc4n2)C(=O)O |
| 3PE2 | 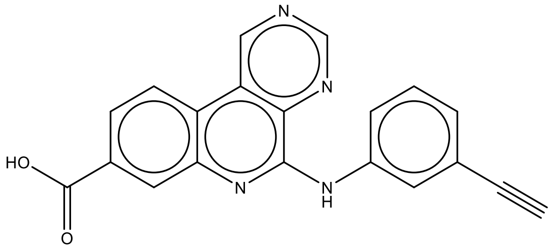<br>C#Cc1cccc(c1)Nc2c3c(cncn3)c4ccc(cc4n2)C(=O)O |
| 3R0T | 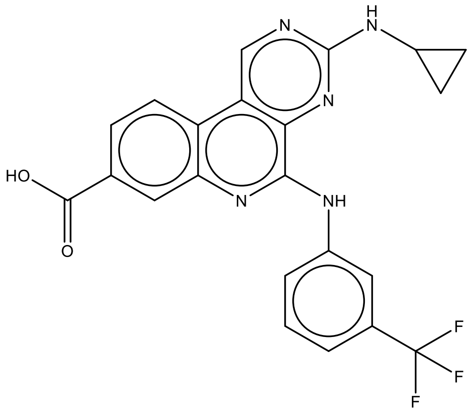<br>c1cc(cc(c1)Nc2c3c(cnc(n3)NC4CC4)c5ccc(cc5n2)C(=O)O)C(F)(F)F |

| | |
|---|---|
| 4RLK | 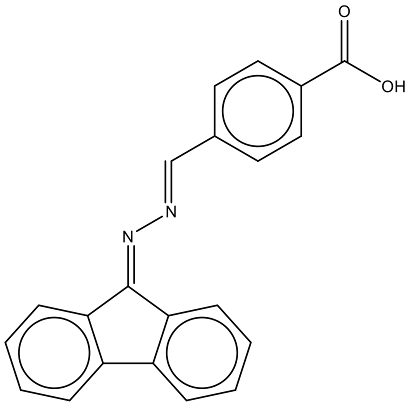<br>c1ccc2c(c1)-c3ccccc3C2=N/N=C/c4ccc(cc4)C(=O)O |
| 4UB7 | 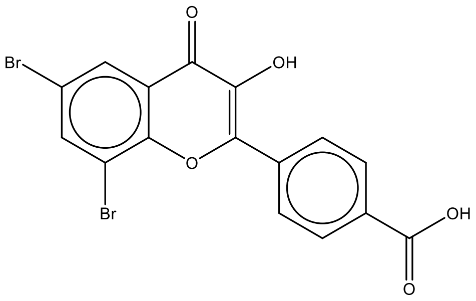<br>c1cc(ccc1C2=C(C(=O)c3cc(cc(c3O2)Br)Br)O)C(=O)O |
| 5B0X | 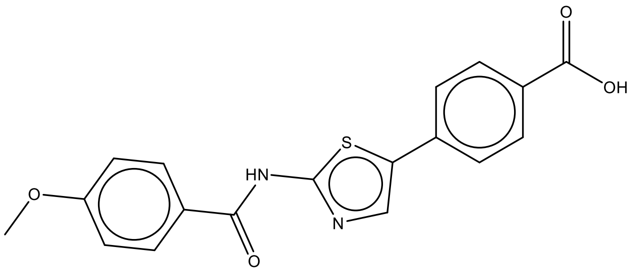<br>COc1ccc(cc1)C(=O)Nc2ncc(s2)c3ccc(cc3)C(=O)O |
| 5M4F | 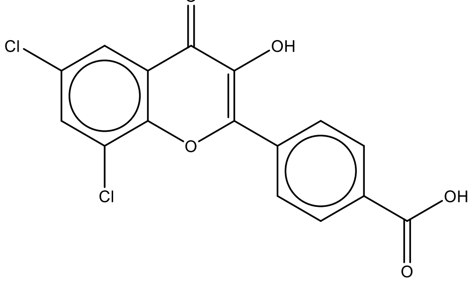<br>c1cc(ccc1C2=C(C(=O)c3cc(cc(c3O2)Cl)Cl)O)C(=O)O |

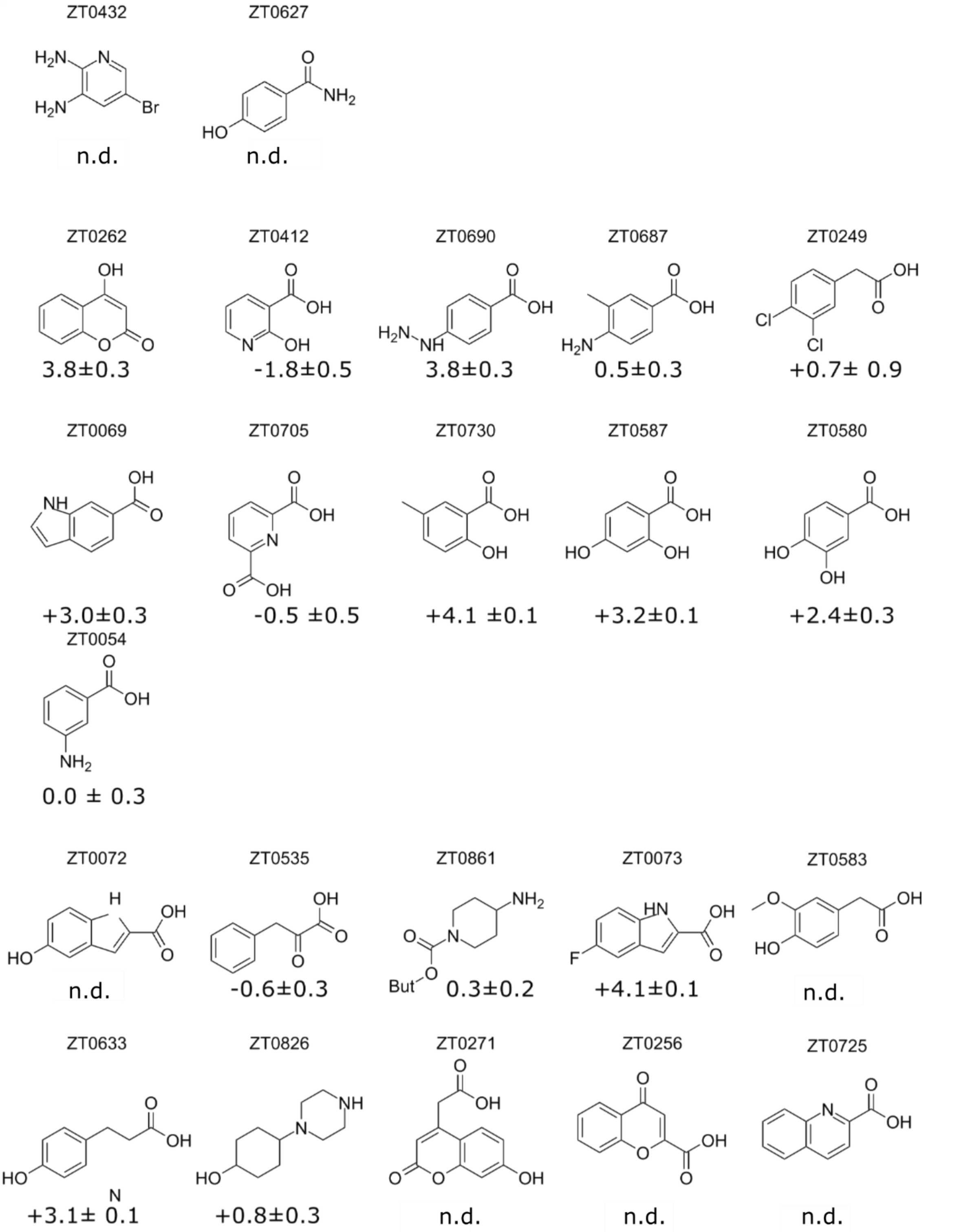


**Supplementary Figure 3:** Structures of 20 fragments identified as binding to the ATP site of CK2α in a recent fragment screen[3]. The thermal shift stabilisation data (°C) are also shown below each structure. Specify units of the thermal shift data.

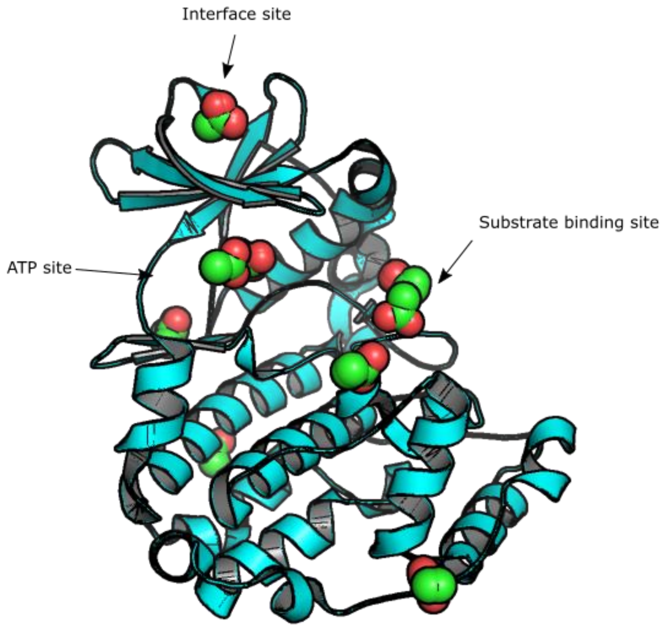


**Supplementary Figure 4:** The various binding sites for acetate identified in CK2α from PDB entries 6Q4Q, 5OSP, 5OSR, 5OSU, 5OTP, 6GMD, 5ORH, 6IGH, 5N9K.

**Supplementary Table 5.** X-ray crystallographic data collection, processing and refinement statistics.

| Description | CK2α in complex with **2** | CK2α in complex with **2** | CK2α in complex with **1** | CK2α in complex with GDP | CK2α in complex with ATP |
|---|---|---|---|---|---|
| PDB code | 6YPG | 6YPH | 6YPJ | 6YPK | 6YPN |
| Ligand | 2 | 2 | 1 | GTP/GDP | ATP/ADP |
| **Data collection and processing:** | | | | | |
| Collection Date | 04/05/2016 | 16/07/2016 | 25/06/2015 | 25/09/2016 | 13/05/2019 |
| Synchrotron | DLS | DLS | DLS | DLS | DLS |
| Beamline | i04 | i04 | I04-1 | i24 | I04-1 |
| X-ray wavelength [Å] | 0.9795 | 0.9795 | 0.9282 | 0.96861 | 0.9159 |
| Space group | $P\ 1\ 2_1\ 1$ | $C\ 2\ 2\ 2_1$ | $P\ 1\ 2_1\ 1$ | $P\ 1\ 2_1\ 1$ | P 1 21 1 |
| Unit cell (a b c) [Å] | 57.65 45.66 63.82 | 64.49 68.95 331.57 | 58.13 46.02 63.71 | 57.70 45.37 63.49 | 57.392 45.225 62.631 |
| (α β γ) [°] | 90.00 111.05 90.00 | 90.00 90.00 90.00 | 90.00 111.13 90.00 | 90.00 110.98 90.00 | 90.00 109.90 90.00 |
| Resolution limits [Å] | 59.56-1.51 (1.55-1.51) | 82.89-1.67 (1.71-1.67) | 54.22-1.64 (1.68-1.64) | 49.70-1.79 (1.84-1.79) | 53.97-1.58 (1.66-1.58) |
| No of reflections, total / unique | 316989 / 47579 | 547559 / 85831 | 123331 / 38677 | 94177 / 28931 | 266973 / 41463 |
| Multiplicity | 6.7 (6.8) | 6.4 (6.2) | 3.2 (2.6) | 3.3 (3.2) | 6.4 (4.5) |
| $R_{merge}$ | 0.055 (1.247) | 0.132 (0.942) | 0.068 (0.643) | 0.138 (0.806) | 0.111 (1.379) |
| $R_{meas}$ | 0.060 (1.348) | 0.144 (1.027) | 0.081 (0.811) | 0.165 (0.970) | 0.120 (1.572) |
| $I/\sigma I$ | 15.5 (1.5) | 6.5 (1.0) | 7.5 (1.3) | 6.5 (1.5) | 8.1 (0.7) |
| $CC_{1/2}$ | 0.998 (0.604) | 0.990 (0.764) | 0.996 (0.575) | 0.982 (0.544) | 0.998 (0.418) |
| Completeness [%] | 97.6 (95.7) | 99.5 (98.3) | 99.9 (99.8) | 99.2 (99.0) | 99.4 (98.6) |
| **Refinement:** | | | | | |
| $R_{work}/R_{free}$ [%] | 23.3 / 26.8 | 25.5 / 27.5 | 19.5 / 23.2 | 17.9 / 21.0 | 20.5/ 24.8 |
| Number of unique / free reflections | 46489 / 2317 | 84814 / 4323 | 38501/ 1963 | 28903 / 1467 | 41367/ 2102 |
| R.m.s deviations: | | | | | |
| Bond lengths [Å] | 0.01 | 0.01 | 0.01 | 0.01 | 0.01 |
| Bond angles [°] | 0.99 | 0.96 | 0.91 | 0.99 | 1.01 |
| Ramachandran analysis: | | | | | |
| Most favoured | 312 ( 96.0%) | 632 ( 97.4%) | 317 ( 97.5%) | 317 ( 97.5%) | 313 ( 96.6%) |
| Allowed | 11 ( 3.4%) | 16 ( 2.5%) | 7 ( 2.2%) | 8 ( 2.5%) | 8 ( 2.5%) |
| Outliers | 2 ( 0.6%) | 1 ( 0.2%) | 1 ( 0.3%) | 0 ( 0.0%) | 3 ( 0.9%) |
| Number of atoms: | | | | | |
| Protein | 2792 | 5517 | 2792 | 2748 | 2789 |
| Solvent atoms | 209 | 314 | 224 | 145 | 191 |
| Heterogen atoms | 239 | 52 | 249 | 28 | 65 |
| Mean/Wilson B-factor [$Å^2$] | 28.29/ 21.56 | 34.90/ 25.34 | 32.33/ 24.47 | 34.21/ 29.95 | 28.83/ 22.29 |

Data for highest resolution shell is shown in parenthesis.

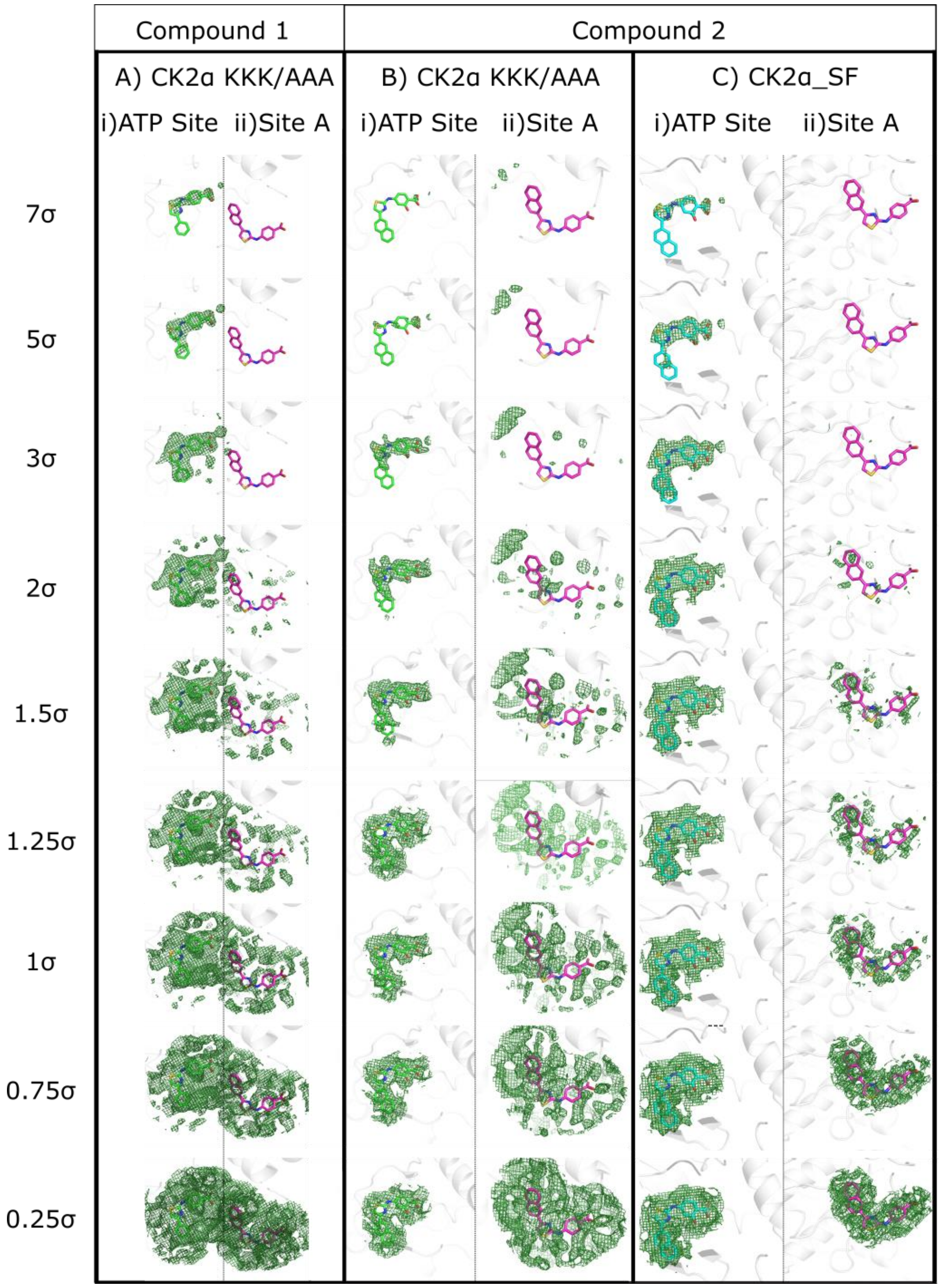


**Supplementary Figure 5.** Electron densities for **1** and **2** before modelling. (A) i) The Fo-Fc map (green) for the ATP site before the ligand has been placed with refined **2** (green) superimposed on the ATP site (PDB:6YPG). ii) The Fo-Fc map (green) for Site A with the predicted binding mode of **2** (purple) is superimposed on the proposed allosteric binding site. (B) The Fo-Fc map (green) for the ATP site before the ligand has been placed with refined **2** (green) superimposed on the ATP site (PDB:6YPH). ii) The Fo-Fc map (green) for Site A with the predicted binding mode of **2** (purple) is superimposed on the proposed allosteric binding site. (C) The Fo-Fc map (green) for the ATP site before the ligand has been placed with refined **1** (blue) superimposed on the ATP site (PDB:6YPH). ii) The Fo-Fc map (green) for Site A with the predicted binding mode of **2** (purple) is superimposed on the proposed allosteric binding site. Contour levels of the maps are shown on the left of the panels.

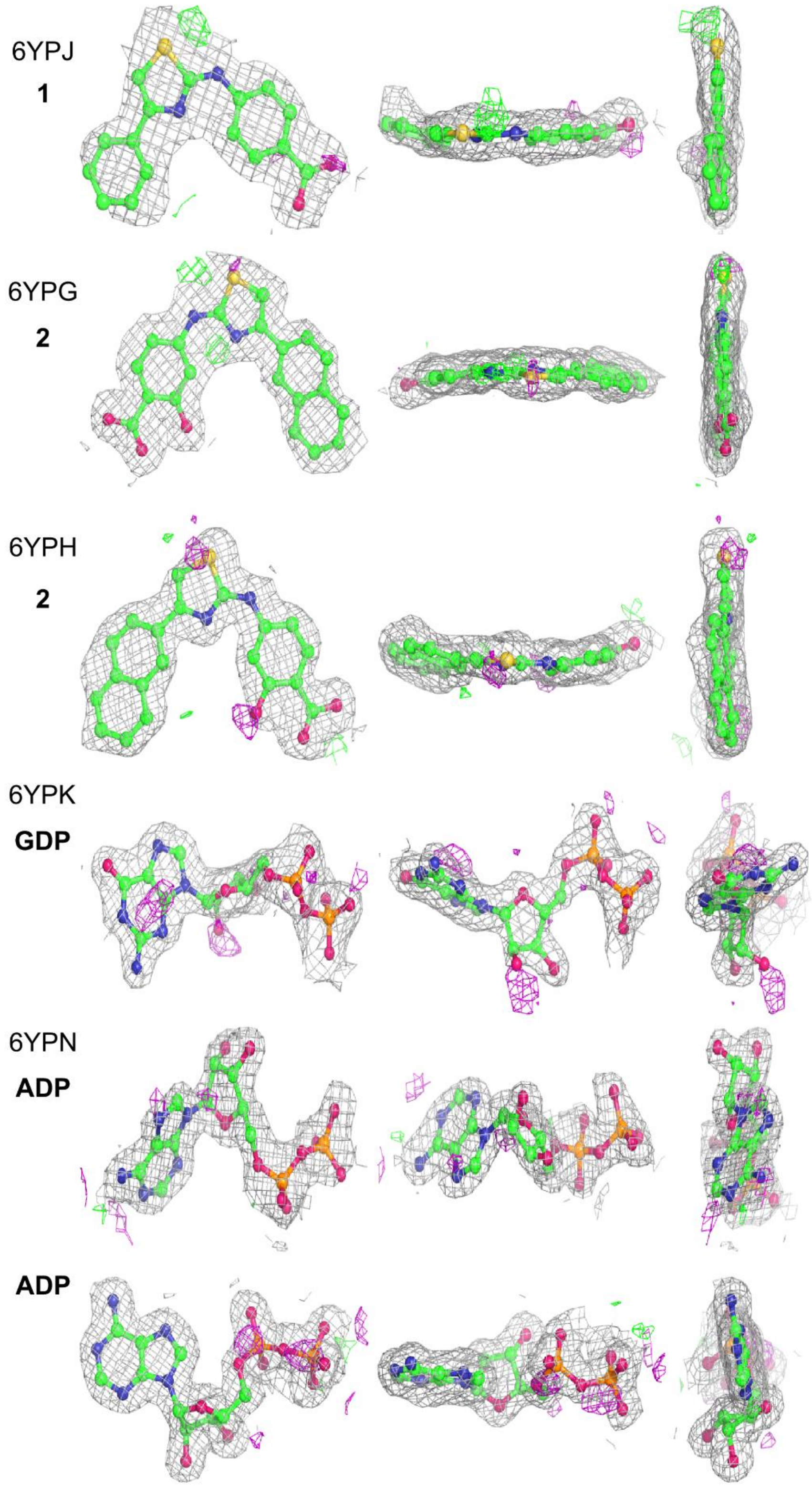


**Supplementary Figure 6.** Ligand electron densities from the PDB validation reports from the deposited structures 6YPG, 6YPH, 6YPJ, 6YPK and 6YPN. The validation reports are calculated by the PDB OneDeposit server (https://deposit-1.wwpdb.org) as part of the PDB deposition process.

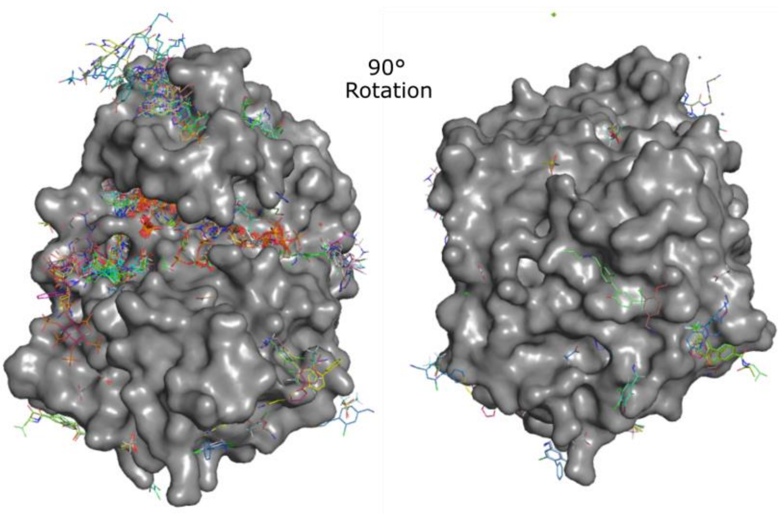


**Supplementary Figure 7**. The binding sites on CK2α that are accessible in the crystal forms used for ligand soaking.

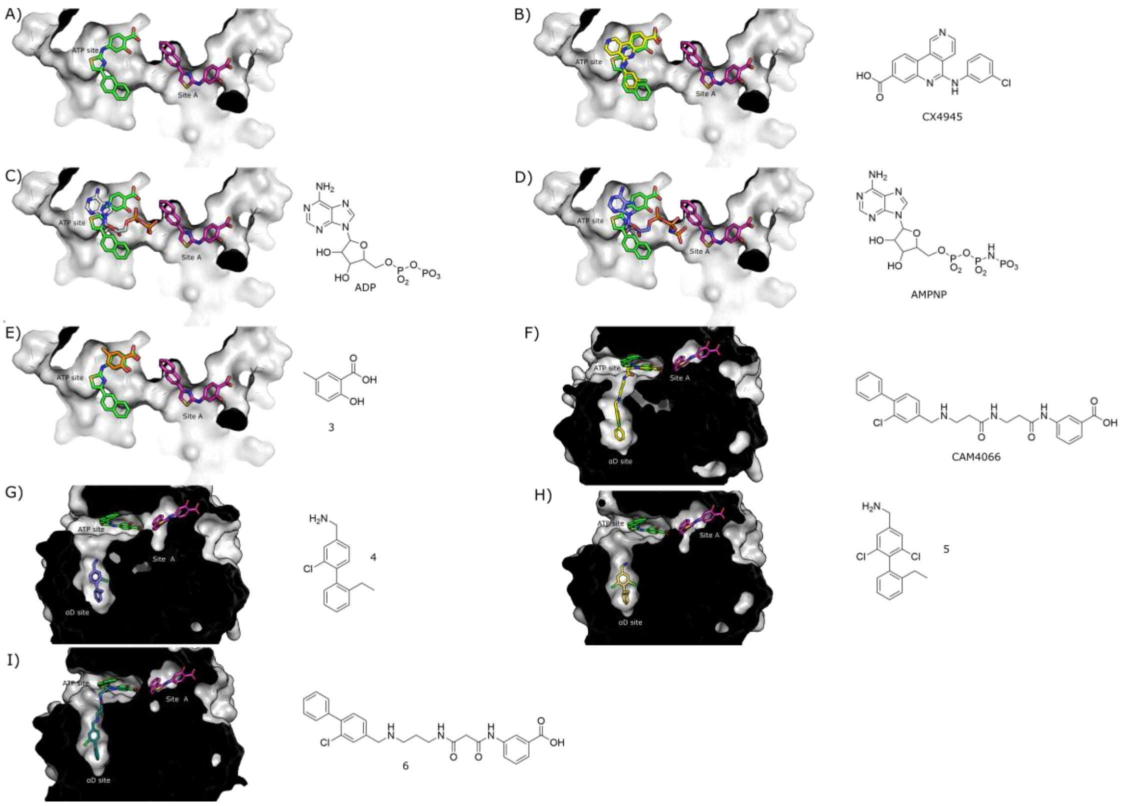


**Supplementary Figure 8**: The overlay of the compounds used as probes in the competition assay with the three sites on CK2α. (A) **2** (green) bound in the ATP site (PDB: 6YPG) with the proposed binding pose of **2** (purple) in site A (as modelled in original publications) These are shown similarly in all other panels, overlaid with: (B) CX4945 (yellow; PDB: 3PE1) bound in the ATP site, (C) ADP (grey; PDB:5ORJ) bound in the ATP site, (D) AMPPNP (blue; PDB:3U87) bound in the ATP site, (E) **3** (orange; PDB:5CSP) bound in the ATP site, (F) **CAM4066** (yellow; PDB:5M08) extending from the αD site to ATP site (G) **4** (blue; PDB:5ORJ) bound the αD site. (H) **4** (yellow; PDB:5OTZ) bound the αD site. (I) **6** (yellow; PDB:5MO8) extending from the αD site to ATP site.

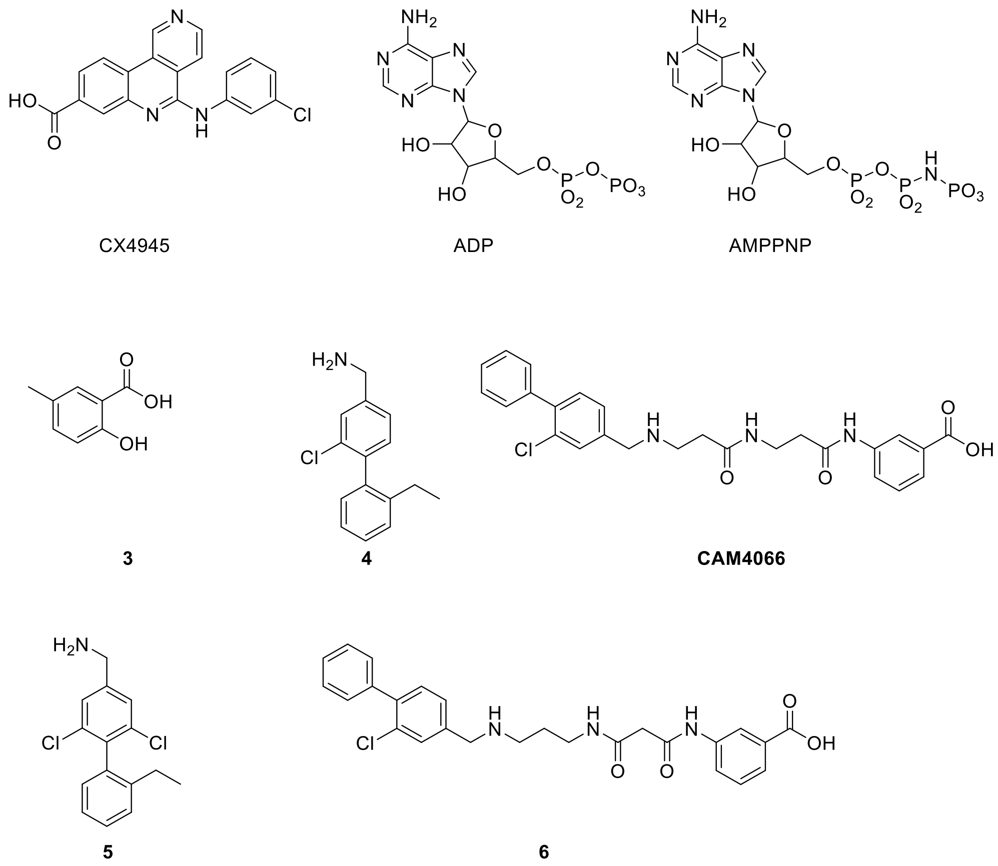


**Supplementary Figure 9:** The six compounds used as probes in HDX and NMR competition studies to determine the binding mode of **1.**

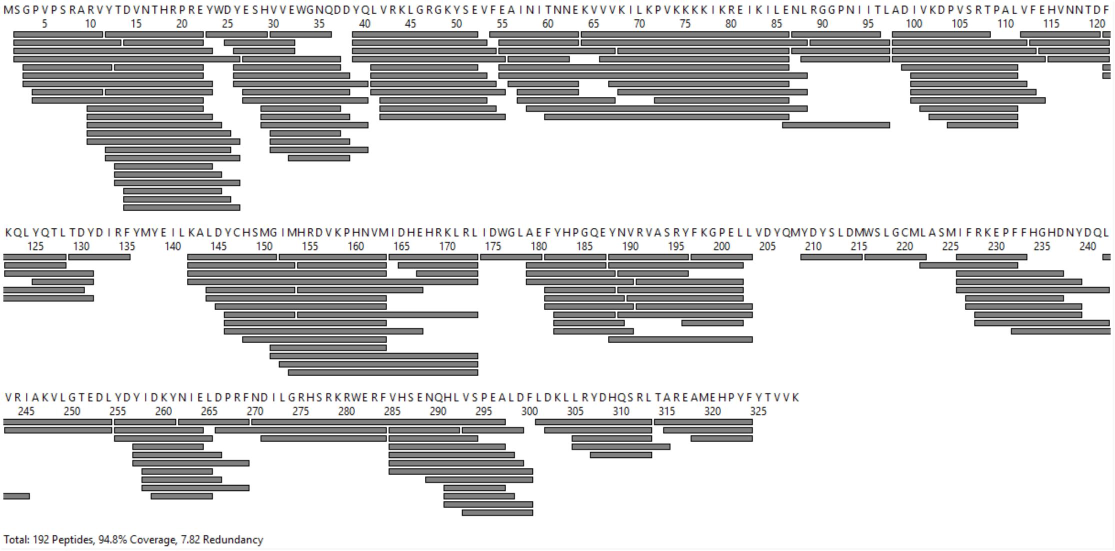


**Supplementary Figure 10.** Coverage map of overlapping peptides recovered in all HDX experiments. Each rectangle represents a peptide recovered with at least one charge state in all HDX experiments and timepoints. There is a total of 192 peptides covering 94.8% of the CK2α sequence with 7.82 redundancy. Regions lacking coverage are 1, 136-141, 204-208, and 325-329.

**Supplementary Table 6. Summary of HDX experimental conditions.**

| Data Set | Experiment A | Experiment B | Experiment C | Experiment D | Shared Across All Experiments |
|---|---|---|---|---|---|
| HDX reaction details | 20 mM Tris<br>500 mM NaCl<br>125 mM $Mg^{2+}$<br>**± 1 mM ADP**<br>$pH_{read}$ = 7.6 | 20 mM Tris<br>500 mM NaCl<br>0.084% MeCN<br>**± 40 µM 6**<br>$pH_{read}$ = 7.6 | 20 mM Tris<br>500 mM NaCl<br>0.16% MeCN<br>**± 180 µM 3**<br>$pH_{read}$ = 7.6 | 20 mM Tris<br>500 mM NaCl<br>0.08% DMSO ±<br>**54 µM 1, 1 mM Mg-AMPPNP, or 37 µM 4**<br>$pH_{read}$ = 7.6 | 20 mM Tris<br>500 mM NaCl<br>$pH_{read}$ = 7.6 |
| HDX time course (s) at 25°C | 10, 100, 1000, 10000, 100000 | | | | 10, 100, 1000, 10000, 100000 at |
| HDX control samples | Unlabeled CK2α | | | | Unlabeled CK2α |
| Back-exchange (mean) | ~30% | | | | ~30% |
| # of peptides | 221 | 213 | 213 | 213 | 192 |
| Sequence coverage | 94.8% | 94.8% | 94.8% | 94.8% | 94.8% |
| Average peptide length / Redundancy | 12.55 / 8.89 | 12.69 / 8.66 | 12.71 / 8.68 | 13.28 / 9.05 | 12.70 / 7.82 |
| Replicates (technical) | 3 | 3 | 3 | 3 | 3 |
| Repeatability (average standard deviation) | 0.058 Da | 0.066 Da | 0.054 Da | 0.058 Da | 0.059 Da |
| Significant difference | ΔHDX greater than 0.25 Da<br>p-value<0.01 | | | | ΔHDX greater than 0.25 Da<br>p-value<0.01 |

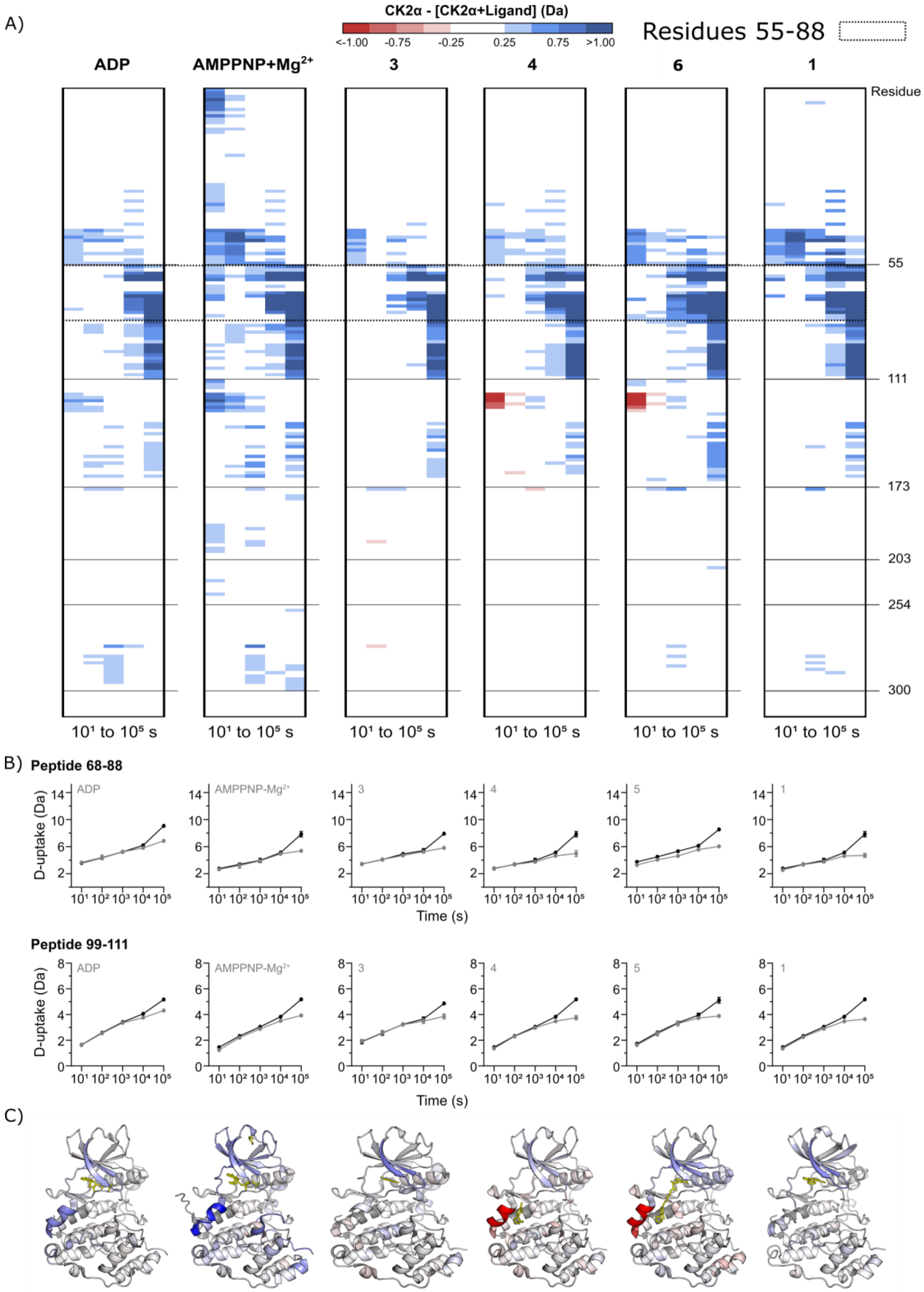


**Supplementary Figure 11. Heat maps showing changes in deuterium uptake upon ligand binding to CK2α.** Heat maps show the change in deuterium uptake upon ligand binding to apo CK2α after $10^1$ s to $10^5$ s deuterium incorporation. All depicted changes are greater than ±0.25 Da that have a p-value less than 0.01 in a Welch's t-test. Labelled residues correspond to the C-terminal boundary of the indicated peptide. Peptides in this map from top to bottom are: 2-11, 2-13, 2-23, 2-26, 3-12, 3-22, 3-23, 4-11, 4-22, 10-22, 10-23, 10-24, 10-25, 10-26, 12-22, 12-23, 12-24, 12-26, 13-22, 13-23, 13-24, 13-26, 14-22, 14-24, 14-25, 14-26, 23-29, 25-32, 26-32, 26-37, 26-38, 26-40, 27-37, 27-38, 27-40, 29-37, 29-38, 29-40, 30-36, 30-37, 30-38, 30-40, 32-38, 39-52, 39-53, 39-54, 39-55, 41-52, 41-53, 41-54, 41-55, 42-53, 42-54, 42-55, 54-63, 55-63, 55-67, 55-86, 55-88, 56-62, 56-63, 57-63, 57-67**, 58-88, 60-86, 64-86, 64-88, 66-86, 67-86, 68-86, 68-88**, 72-86, 86-97, 87-96, 87-97, 88-97, 89-97, 98-108, 98-112, 98-113, 98-114, 99-111, 100-111, 100-112, 100-113, 100-114, 101-111, 102-111, 104-111, 112-120, 113-121, 114-121, 115-121, 121-128, 121-130, 121-131, 122-128, 122-131, 125-131, 129-135, 142-151, 142-153, 142-163, 142-173, 144-153, 144-163, 145-163, 146-153, 146-163, 146-167, 148-163, 151-163, 151-173, 152-163, 152-173, 153-173, 154-163, 154-167, 154-173, 164-173, 165-173, 167-173, 174-180, 179-187, 179-188, 179-190, 181-187, 181-188, 181-189, 181-190, 182-188, 182-189, 182-190, 188-196, 188-202, 188-203, 189-196, 189-202, 189-203, 190-202, 191-202, 191-203, 196-202, 197-203, 209-215, 216-222, 222-232, 226-233, 226-237, 226-239, 226-242, 227-237, 227-239, 228-239, 228-242, 232-245, 242-254, 243-254, 255-261, 255-264, 255-265, 257-264, 257-266, 257-269, 258-265, 258-266, 258-269, 259-265, 262-269, 266-284, 270-297, 271-284, 285-292, 285-294, 285-297, 285-298, 285-299, 285-300, 289-300, 291-297, 291-298, 291-300, 293-299, 293-300, 301-313, 302-313, 305-313, 305-315, 307-313, 314-324, 315-324, 318-324.

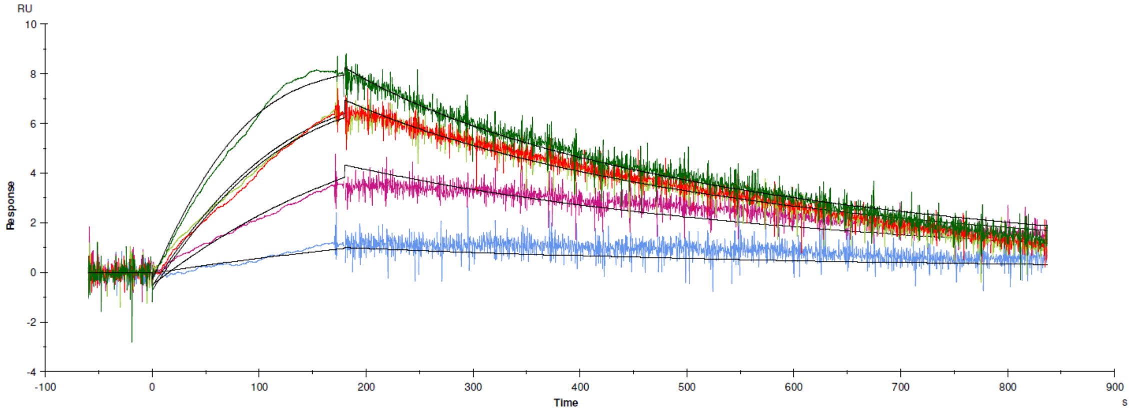


**Supplementary Figure 12. Kinetics of CX4945 binding to CK2α.** Binding of **CX4945** to CK2α as determined by SPR. $k_{on}$ = 1.500 × $10^6$ $M^{-1}s^{-1}$, $k_{off}$ = 0.003897 $s^{-1}$, $K_D$ = 2.6 nM. **CX4945** concentrations were 15 nM (green), 10 nM (orange), 5 nM (purple) and 1 nM (blue).

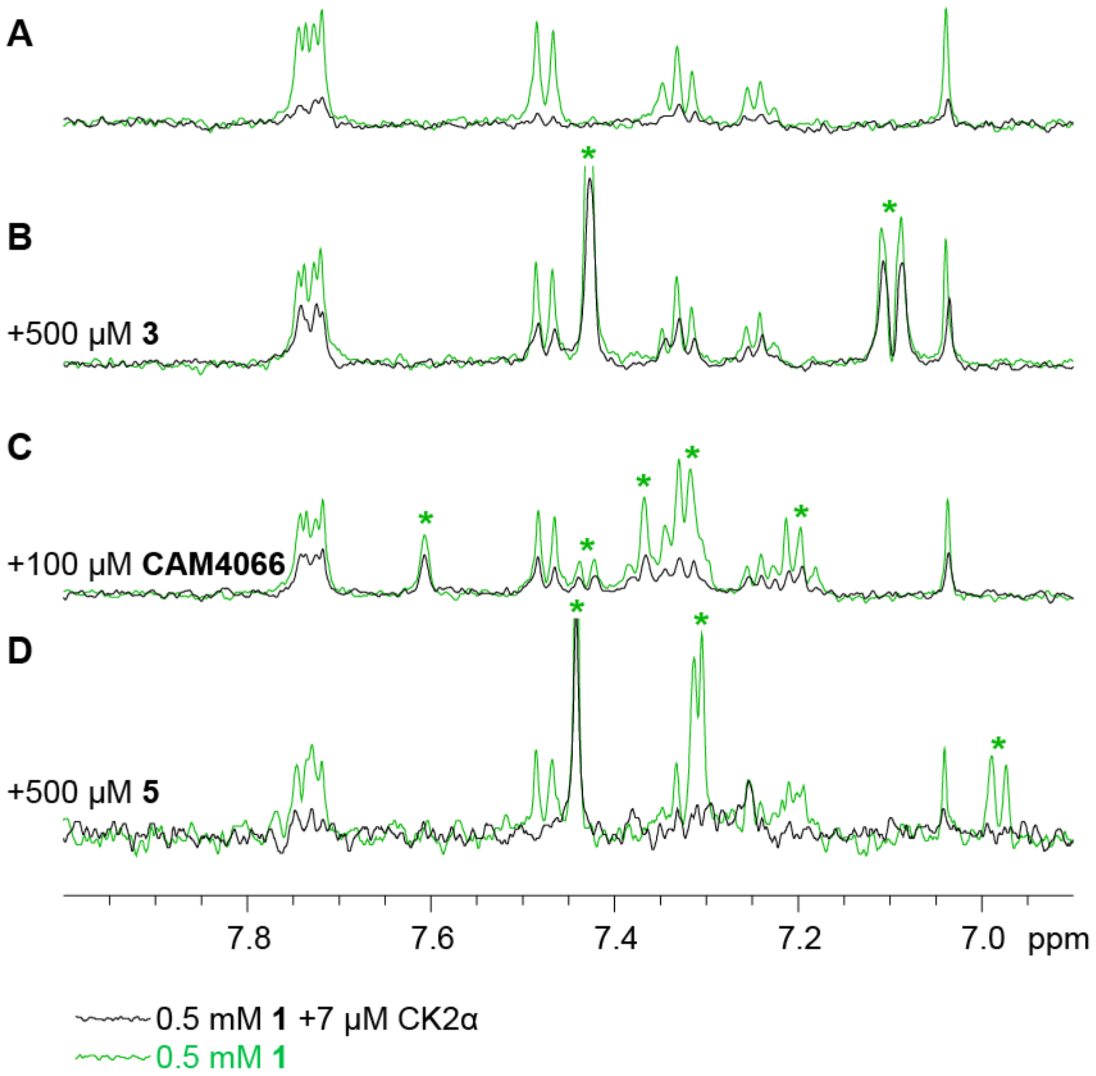


**Supplementary Figure 13.** Relaxation-edited (CPMG) $^1$H NMR analysis of the binding of 1 to CK2α in competition with various ligands. All panels are overlays of spectra recorded in the presence (black) and absence (green) of CK2α. CPMG spectra are of (A) **1**, (B) **1** and **3**, (C) **1** and **CAM4066**, and (D) **1** and **5.** Peaks arising from competitors are indicated by a *.

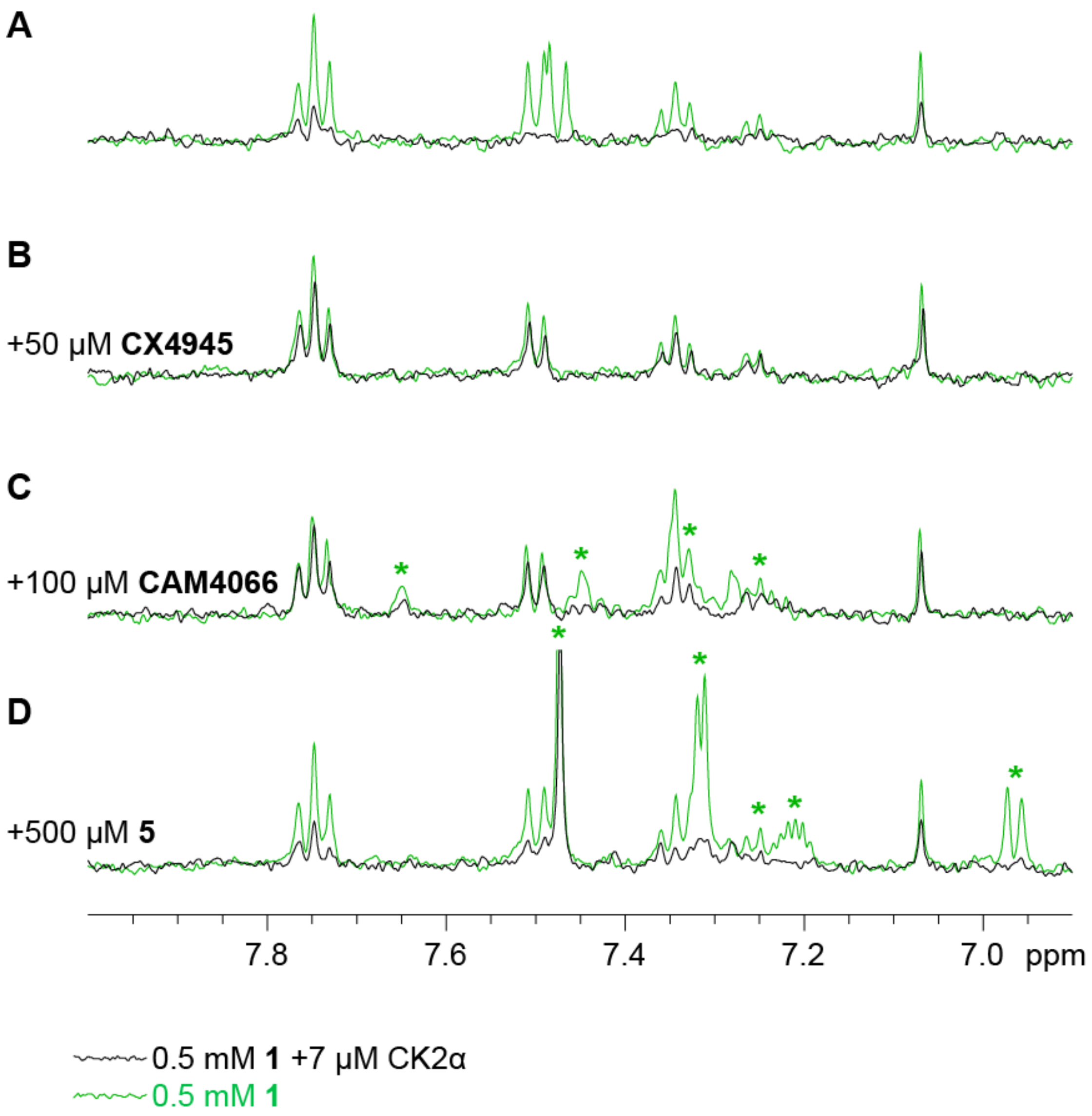


**Supplementary Figure 14**. Relaxation-edited (CPMG) [1]H NMR analysis of the binding of **1** to CK2$\alpha$ KKK/AAA mutant in competition with various ligands. All panels are overlays of spectra recorded in the presence (black) and absence (green) of CK2$\alpha$. CPMG spectra are of (A) **1**, (B) **1** and **CX4945**, (C) **1** and **CAM4066**, and (D) **1** and **5.** Peaks arising from competitors are indicated by a *.

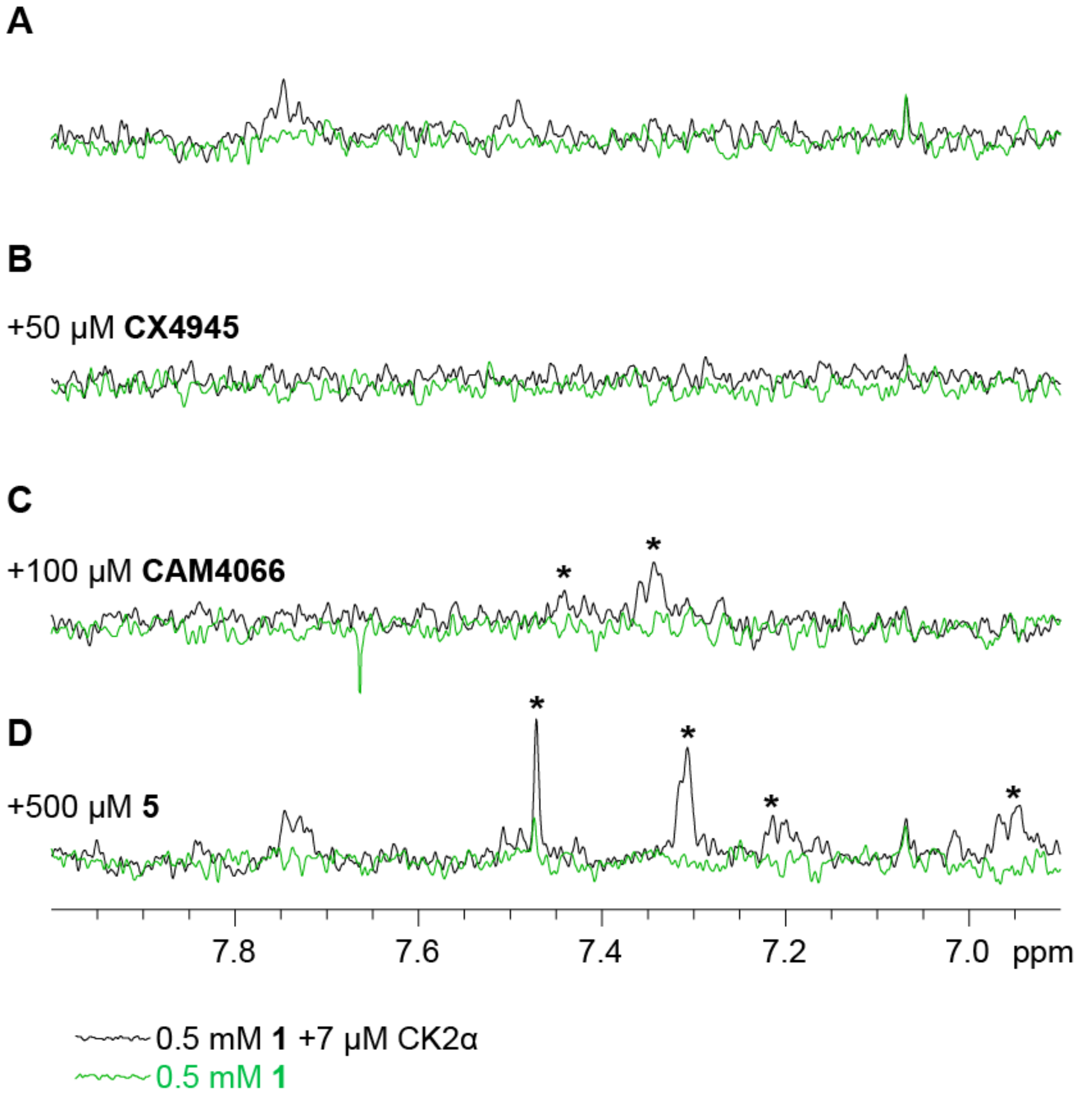


**Supplementary Figure 15. Saturation Transfer Difference (STD) $^{1}$H NMR analysis of the binding of 1 to the CK2α KKK/AAA mutant in competition with various ligands.** All panels are overlays of spectra recorded in the presence (black) and absence (green) of CK2α. STD spectra are of (A) **1**, (B) **1** and **CX4945**, (C) **1** and **CAM4066**, and (D) **1** and **5.** Peaks arising from competitors are indicated by a *.

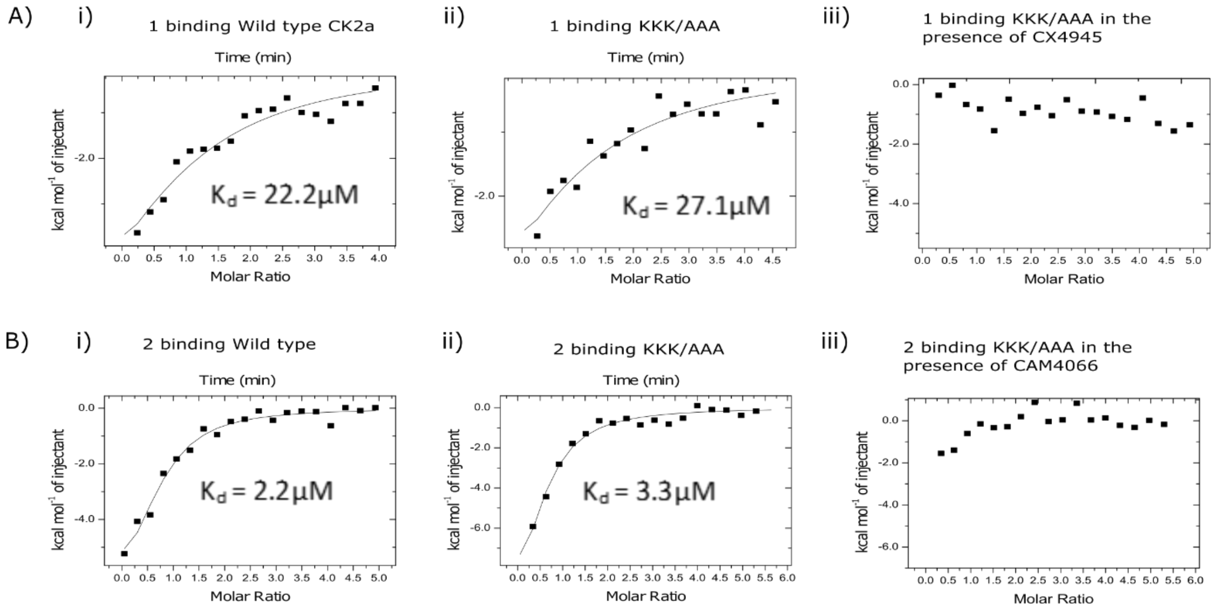


**Supplementary Figure 16**: A) ITC traces of titration of **1** into (i) CK2α WT (as in Figure 5), (ii) the KKK/AAA mutant and (iii) the KKK/AAA mutant in the presence of 100 μM **CX4945**. B) ITC traces of titration of **2** into (i) CK2α WT (as in Figure 5), (ii) the KKK/AAA mutant and (iii) the KKK/AAA mutant in the presence of 100 μM **CX4945**.

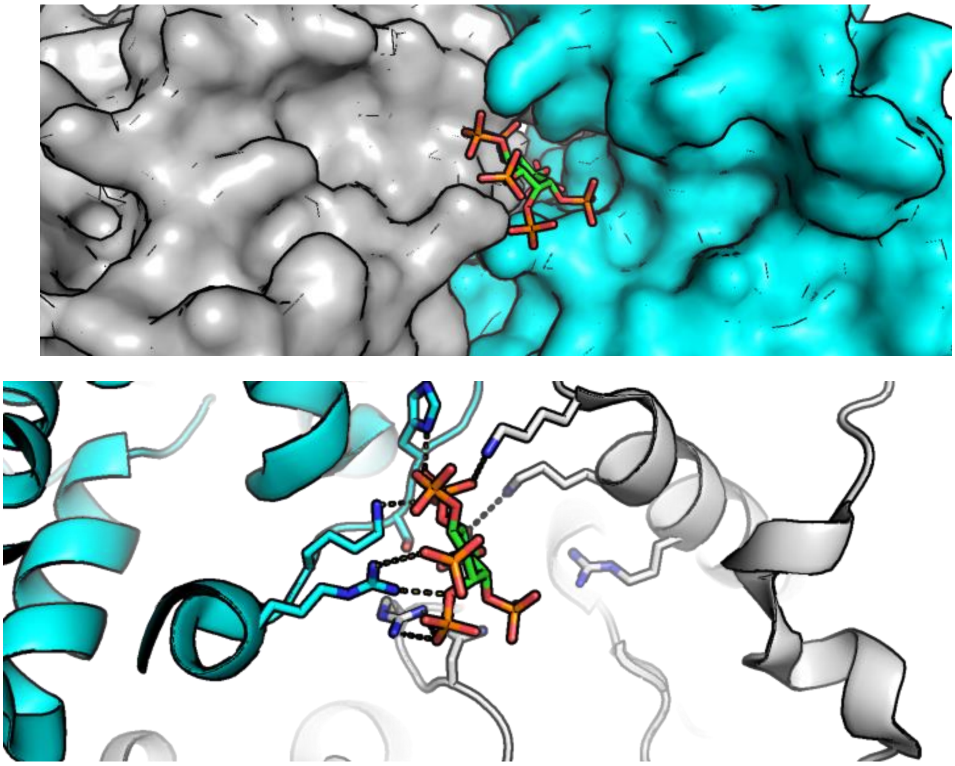

**Supplementary Figure 17:** The binding site of IP6 for CK2α is partially formed by crystal contacts. The binding site of IP6 presented in 3W8L[4] is clearly composed of residues from two separate monomers (blue and grey) of CK2α. As CK2α does not form a dimer in solution it is unlikely that this is a relevant site.

### Materials and Methods

#### Modelling

Modelling of binding of **1** and **2** to  CK2α was done using the GOLD software package (CCDC) using the structure from PDB entry 3PE1 as the target. The site of interest was defined as a 20 Angstrom sphere around the ligand CX4945, which includes both the ATP site and the proposed allosteric site. Ligand files were generated with grade (Global Phasing). The conserved water interacting with Lys68 was included but no other waters. The ChemPLP scoring system (CCDC) was used to rank the results. Coordinates of the modelling results are included as supplementary files.

#### Expression and purification

Expression and purification of CK2α (wild type, KKK/AAA and SF) was done as described before[5]. For the surface plasmon resonance experiments a new construct carrying two His-tags, one at the N-terminus and one at the C-terminus was created. CK2α was amplified by PCR using primers “AGCTA GAATTCGGAGAAGGCGAGGGTAGTGAAGGCCCCGTGCCAAGCAGGGCCAGAGT” and “ATACTCAAGCTTAATGGTGATGGTGATGGTGGAACCCTGGAAGTACAGGTTTTCCTTCACAACAGTGTA GAAATAGGGGTG”. The resulting fragment was digested with *Eco*RI and *Hind*III restriction enzymes and ligated into a pHAT4 vector digested with the same enzymes. The sequence of the protein that the resulting construct (pHAT4_CK2-TEV-L-His) encodes for is shown below with His-tags (bold) and two TEV cleavage sites (underlined).

MNTI**HHHHHH**NTSGSGGGGGRLVPRGSMSENLYFQGSMDIEFGEGEGSEGPVPSRARVYTDVNTHRPSEYW<br>
DYESHVVEWGNQDDYQLVRKLGRGKYSEVFEAINITNNEKVVVKILKPVKKKKIKREIKILENLRGGPNII<br>
TLADIVKDPVSRTPALVFEHVNNTDFKQLYQTLTDYDIRFYMYEILKALDYCHSMGIMHRDVKPHNVMIDH<br>
EHRKLRLIDWGLAEFYHPGQEYNVRVASRYFKGPELLVDYQMYDYSLDMWSLGCMLASMIFRKEPFFHGHD<br>
NYDQLVRIAKVLGTEDLYDYIDKYNIELDPRFNDILGRHSRKRWERFVHSENQHLVSPEALDFLDKLLRYD<br>
HQSRLTAREAMEHPYFYTVVKENLYFQGSSGSGSGSS**HHHHHH***

Expression of this protein was done as with other CK2α construct, using the BL21(DE3) strain of *E. coli.*  Single colonies of the cells were grown in 6 l 2xTY with 100 µg/mL ampicillin at 37 °C until $OD_{600}$ of 0.6 after which 0.4 mM IPTG was used to induce expression. After three hours of culturing at 37 °C the cell were harvested by centrifugation at 4,000 g for 20 minutes. The cells were resuspended in 20 mM Tris pH 8.0, 500 mM NaCl and lysed using an Emulsiflex C5 homogeniser. DNaseI was added to aid with digestion of *E. coli* genomic DNA and protease inhibitor tablets we are added to minimise proteolytic degradation. Following centrifugation for 30 min at 15,000 *g*, the clarified supernatant was loaded onto 4 ml $Ni^{2+}$-NTA agarose column and washed with the lysis buffer, followed by 20 mM Tris pH 8.0, 500 mM NaCl, 20 mM imidazole and the protein eluted with 20 mM Tris pH 8.0, 500 mM NaCl, 300 mM imidazole. Peak fractions, as analysed by SDS-PAGE, were pooled together and dialysed overnight against 4 l of 20mM Tris pH 8.0, 500 mM NaCl at +4°C. The dialysed sample was diluted to ~65 mM salt concentration in two batches and immediately loaded onto HiTrapQ HP column to minimise aggregation in low ionic strength. CK2α was eluted using a linear salt gradient. Peak fractions of each of the two chromatographic steps were pooled, concentrated to ca. 5 ml in volume and purified with size exclusion chromatography on a Superdex 75 16/60 HiLoad column using 20 mM Tris pH 8.0, 500 mM NaCl buffer. Peak fractions were pooled, concentrated to ca. 10 mg/ml using Vivaspin 20 5000 MWCO centrifugal concentrators and flash frozen in small aliquots.

#### X-ray crystallography

Crystallisation, soaking of ligands and structure determination was done as described before[5].  X-ray diffraction data were collected at the Diamond Light Source and data from automated data processing with autoProc were used for the structure determination. Fo-Fc electron density maps for all the ligands, calculated prior to ligand placement in the model, are shown in Supplementary Figure

5 and the final densities, as calculated by PDB AutoDep software, are shown in Supplementary Figure 6. All coordinates have been deposited to the Protein Data Bank under accession numbers 6YPG, 6YPH, 6YPJ, 6YPK, 6YPN. We will release the atomic coordinates and experimental data upon article publication. Data collection and refinement statistics are shown in Supplementary Table 5.

#### Hydrogen-deuterium exchange mass spectrometry

Stocks of 25 µM CK2α with and without ligand were prepared in 20 mM Tris-HCl, 500 mM NaCl, pH 8.0 at 25°C. In some experiments a small percentage of organic solvent was added to increase solubility. Variations and ligand concentrations are noted in Supplementary Table 6. Stocks were equilibrated at 4°C for 10 minutes and then diluted 1:24 with the same buffer containing $H_2O$ for controls ($pH_{read}$=8.0 at 25°C) or $D_2O$ (final $D_2O$ of 96% (v/v) with $pH_{read}$=7.6 at 25°C). After 10 s, $10^2$ s, $10^3$ s, $10^4$ s, and $10^5$ s, samples were mixed 1:1 with 0.8 % formic acid, 1.6 M guanidinium chloride, pH 2.3 to give a final pH of 2.6 at 0 °C, and flash frozen in liquid nitrogen for storage at –80°C. Samples were thawed and LC-MS performed using a Waters HDX manager and SYNAPT G2-Si Q-Tof. Three technical replicates of each sample were analyzed in a random order. Samples were digested online by *Sus scrofa* Pepsin A (Waters Enzymate BEH) at 15°C and peptides trapped on a C18 pre-column (ACQUITY UPLC Peptide BEH C18 VanGuard Pre-column) for 3 min at 100 µL/min and 1°C. The LC buffer was 0.1% formic acid. Peptides were separated over a C18 column (Waters acquity UPLC BEH) and eluted with a linear 3-40% (v/v) acetonitrile gradient for 7 min at 40 µL/min and 1°C. Raw data of control samples were processed by PLGS (Waters Protein Lynx Global Server 3.0.2) using a database containing *Sus scrofa* Pepsin A and *Homo sapiens* CK2α. In PLGS, the minimum fragment ion matches per peptide was 3, and methionine oxidation was allowed. The low and elevated energy thresholds were 250 and 50 counts respectively, and overall intensity threshold was 750 counts. Labelled data were analyzed in DynamX 3.0 with peptide filters of 0.3 products per amino acid and 1 consecutive product. All mass spectrometry data were acquired using positive ion mode in either HDMS or HDMSE modes. HDMSE mode was used to collect both low (6V) and high (ramping 22-44 V) energy data-independent peptide fragmentation data for peptide identification. HDMS mode was used to collect low energy ion data for all deuterated samples. All samples were acquired in resolution mode. Capillary voltage was set to 2.8 kV for the sample sprayer. Desolvation gas was set to 650 L/hour at 175°C. The source temperature was set to 80°C. Cone and nebulizer gas was flowed at 90 L/hour and 6.5 bar, respectively. The sampling cone and source offset were both set to 30 V. Data were acquired at a scan time of 0.4 s with a range of 100-2000 m/z. Mass correction was done using [Glu1]-fibrinopeptide B as a reference mass. To allow access to the HDX data of this study, the HDX data summary table (Supplementary Table 6) and the HDX data table (Supplementary Table X) are included per consensus guidelines[6].

#### Ligand-observe NMR experiments

In the control experiment a stock solution of 2.5% DMSO-$d_6$, 7 µM CK2α WT in 0.5mM Tris, 200 mM NaCl, pH 8.0 was prepared. This was split into two samples. Ligand in DMSO-$d_6$ or DMSO-$d_6$ alone was added to each such that one sample contained 5% DMSO-$d_6$ with a total volume of 180 µL, and the second sample contained 5% DMSO-$d_6$ and 500 µM **1**. In a typical competition experiment a stock solution of 50-500µM competing ligand, 2.5% DMSO-$d_6$, 7 µM CK2α WT in 0.2mM Tris, 100 mM NaCl, pH 8.0 was prepared. This was again split into two samples. One sample contained 5% DMSO-$d_6$ and 50-500 µM competing ligand with a total volume of 180 µL. The second sample contained 5% DMSO-$d_6$, 500 µM **1** and 50-500 µM competing ligand.

NMR data were recorded at 300 K with a Bruker AVANCE spectrometer operating at 500 MHz and equipped with a room-temperature probe. CPMG-filtered 1D $^1$H experiments [7] were run with 600 loops of length 1 ms. Water suppression was achieved with low-power presaturation and DPFGSE

WATERGATE W5 [8,9]. Saturation transfer difference (STD) experiments [10] were acquired with 2 s of alternating on- and off-resonance saturation (20 ms Gaussian pulse, +0.8 and –5 ppm), a $T_{1\rho}$ filter to suppress protein signals, and DPFGSE WATERGATE W5. WATER-LOGSY experiments [11] were recorded using a 20 ms Gaussian to invert the water resonance, a mixing time of 1.2 s, a $T_2$ filter to suppress protein signals, and DPFGSE WATERGATE W5.

#### Isothermal calorimetry

ITC All ITC experiments were performed at 25 °C using a MicroCal iTC200 instrument (GE Healthcare). CK2αWT (20 mg/mL, 20 mM Tris pH 8.0, 500 mM NaCl) was diluted in Tris buffer (200 mM Tris, 300 mM NaCl, 10% DMSO) and concentrated to 20-50 μM. Compounds in 100x stock solutions were diluted into the buffer ensuring that the DMSO concentrations were carefully matched. In a typical experiment CK2αWT (10 μM) was loaded into the sample cell and 0.4-2.0 mM of the ligand was titrated in eighteen 2 μL injections of 2 s duration at 150 s intervals, stirring at 750 rpm. Heats of dilution were determined in identical experiments, but without protein in the cell. The data fitting was performed with a single site binding model using the Origin software package.

#### Surface Plasmon Resonance

An Ni-NTA sensor chip (GE Healthcare) was activated with 0.5 mM $NiCl_2$, 0.2 μM CK2α with both N- and C-terminal His-tags was immobilised onto the activated Ni-NTA sensor chip in 20 mM Tris pH 8.0, 500 mM NaCl. Binding of **CX4945** was determined in 0.01 M HEPES pH 7.4, 0.15 M NaCl, 0.05% v/v TWEEN 20, 0.5% DMSO at five different concentrations (15 nM, 10 nM, 5 nM, 1 nM and 0 nM). The chip surface was regenerated with an initial wash of 500 mM EDTA followed by 4 M guanidinium chloride, 250 mM EDTA, 50 mM NaOH. The reference channel was activated with 0.5 mM $NiCl_2$. All data were processed using the manufacturer's BIAevaluation software.


1. Bestgen, B. *et al.* 2-Aminothiazole Derivatives as Selective Allosteric Modulators of the Protein Kinase CK2. 2. Structure-Based Optimization and Investigation of Effects Specific to the Allosteric Mode of Action. *J. Med. Chem.* **62**, 1817–1836 (2019).
2. Battistutta, R. *et al.* Unprecedented selectivity and structural determinants of a new class of protein kinase CK2 inhibitors in clinical trials for the treatment of cancer. *Biochemistry* **50**, 8478–8488 (2011).
3. De Fusco, C. *et al.* A fragment-based approach leading to the discovery of a novel binding site and the selective CK2 inhibitor CAM4066. *Bioorg. Med. Chem.* **25**, 3471–3482 (2017).
4. Lee, W.-K. *et al.* Structural and functional insights into the regulation mechanism of CK2 by IP6 and the intrinsically disordered protein Nopp140. *Proc.Natl.Acad.Sci.USA* **110**, 19360–19365 (2013).
5. Brear, P. *et al.* Specific inhibition of CK2 alpha from an anchor outside the active site. *Chem Sci* **7**, 6839–6845 (2016).
6. Masson, G. R. *et al.* Recommendations for performing, interpreting and reporting hydrogen deuterium exchange mass spectrometry (HDX-MS) experiments. *Nat. Methods* **16**, 595–602 (2019).
7. Hajduk, P. J., Olejniczak, E. T. & Fesik, S. W. One-dimensional relaxation- and diffusion-edited NMR methods for screening compounds that bind to macromolecules. *J. Am. Chem. Soc.* **119**, 12257–12261 (1997).
8. Stott, K., Keeler, J., Hwang, T. L., Shaka, A. J. & Stonehouse, J. Excitation Sculpting in High-

Resolution Nuclear Magnetic Resonance Spectroscopy: Application to Selective NOE Experiments. *J. Am. Chem. Soc.* **117**, 4199–4200 (1995).

9. Liu, M. *et al.* Improved Watergate Pulse Sequences for Solvent Suppression in NMR Spectroscopy. *J. Magn. Reson.* **132**, 125–129 (1998).

10. Mayer, M. & Meyer, B. Characterization of ligand binding by saturation transfer difference NMR spectroscopy. *Angew. Chemie - Int. Ed.* **38**, 1784–1788 (1999).

11. Dalvit, C., Fogliatto, G., Stewart, a, Veronesi, M. & Stockman, B. WaterLOGSY as a method for primary NMR screening: practical aspects and range of applicability. *J. Biomol. NMR* **21**, 349–59 (2001).
